# Cell-cycle-resolved Hi-C reveals unexpected plasticity of A/B compartments across interphase

**DOI:** 10.64898/2025.12.06.692720

**Authors:** Linda Choubani, Hisashi Miura, Takako Ichinose, Asami Oji, Saori Takahashi, Rory T. Cerbus, Ichiro Hiratani

## Abstract

The spatial organization of chromatin into active (A) and inactive (B) nuclear compartments is fundamental to genome regulation, yet their cell-cycle dynamics remain largely unexplored. Most research on chromatin dynamics during the cell cycle has primarily focused on events surrounding mitosis, providing only limited insight into chromatin behavior during S-phase. To address this gap, we developed a simple, drug-free approach that combines the Fucci cell-cycle indicator with *in situ* Hi-C to comprehensively analyze A/B compartment dynamics throughout interphase in mouse embryonic stem cells (mESCs). Unexpectedly, and contrary to prevailing views, we found that A/B compartment strength increased abruptly upon S-phase entry, stabilized during S-phase, and subsequently declined in late S/G2. This abrupt strengthening, which we termed ‘compartment maturation,’ required passage through the G1/S transition but was independent of active DNA synthesis. This maturation involved substantial architectural remodeling, particularly within the A compartment, which consolidated into a more organized structure as individual A domains rearranged to form long-range interactions. Moreover, compartment maturation was not limited to mESCs but was also evident across different developmental contexts in mice. Based on these observations, we propose a revised, stepwise model of nuclear compartmentalization during cell-cycle progression, consisting of four distinct stages: chromosome unfolding (G1), chromatin maturation (G1/S), stabilization (S phase), and refolding (G2). These findings reveal the unexpected plasticity of A/B compartments and underscore the G1/S transition as a critical period for their reorganization.

## Introduction

The three-dimensional (3D) organization of the genome within the nucleus plays a fundamental role in regulating essential cellular processes, including gene expression, DNA replication, and repair^1,2^. High-throughput chromosome conformation capture (Hi-C) techniques^3^ have revolutionized our ability to probe this organization, revealing the existence of megabase (Mb)-scale chromosomal compartments in interphase cells. These appear as a characteristic plaid pattern on Hi-C contact maps, reflecting the segregation of chromatin into self-interacting active (A) and inactive (B) nuclear compartments^3^. This pattern is identified computationally through principal component analysis (PCA) of the Hi-C contact matrix, where the first principal component (PC1) serves as a quantitative eigenvector; genomic bins are assigned to A (positive PC1 values) or B (negative PC1 values) compartments based on their correlated interaction profiles^3^. Broadly speaking, A and B compartments correspond to euchromatin and heterochromatin, respectively, in the classical cytological sense, and are typically located in the nuclear interior and at the nuclear periphery^4^.

Despite their fundamental significance in genome function, the principles governing compartment formation remain largely elusive. Over the past decade, researchers have increasingly focused on the cell cycle regulation to unravel the principles of chromatin organization, given that compartments are largely lost during mitosis and re-established during interphase^5^. This process begins as chromosomes compact into their distinct, rod-like shapes during prophase^6,7^. Following mitosis, as cells traverse the M/G1 transition, chromosomes expand from this highly condensed state and gradually unfold to re-establish interphase chromatin architecture. Time-course Hi-C studies of this critical window^8–11^ supported a “compartment expansion” model, positing that compartment boundaries are established soon after mitotic exit, followed by a gradual strengthening of chromatin interactions and their simultaneous expansion outward from the diagonal on Hi-C contact maps, reflecting the progressive increase in long-range interactions during G1. This progression culminates in the model proposed in the pioneering study by Nagano *et al.*^12^, where compartmentalization progressively strengthens after mitotic exit, peaking in G2 just before its rapid collapse in the subsequent mitosis.

Despite these insights, significant gaps remain in our knowledge. Although compartments were among the first structures identified in Hi-C maps, they have often not been the primary focus of studies and are frequently treated as binary, static entities. Even when they have been investigated, the emphasis has typically been placed on Hi-C PC1-defined compartment boundaries rather than on the internal dynamics of the compartments themselves. Furthermore, variability in compartment calculation methods across studies complicates direct comparisons and hampers a comprehensive understanding. Importantly, while previous studies have illuminated events around mitosis, they have left the S-phase notably underexplored. This is a critical gap given the strong correlation between A/B compartments and DNA replication timing, whereby A regions replicate in early S-phase and B regions in late S-phase^13,14^. The fragmented nature of the existing cell-cycle Hi-C datasets is further complicated by the reliance on synchronization drugs in most studies, which can introduce population heterogeneity and confound biological interpretation^15^.

In this study, we address these challenges by thoroughly investigating the temporal dynamics of interphase A/B compartments, with a special emphasis on the G1-to-S transition, a crucial yet understudied period for the coordinated establishment of 3D genome structure and DNA replication.

## Results

### Drug-free cell cycle-phasing system in mESCs

To investigate how A/B nuclear compartments change throughout the cell cycle, we established a drug-free synchronization approach in mouse embryonic stem cells (mESCs) to avoid the heterogeneity associated with the use of chemical inhibitors. For high-resolution cell-cycle discrimination, we used the Fucci (SA) mESC line^16,17^, which expresses Cdt1-mCherry in G1 (degraded at the G1/S transition) and Geminin-mVenus in S/G2/M (accumulating from S-phase entry onwards) (**Fig. 1A)**. The reciprocal fluorescence patterns of Cdt1 and Geminin enabled distinct and dynamic visualization of cell cycle phases, thereby allowing accurate sorting of highly pure cell populations for Hi-C without relying on chemical synchronization.

**Figure 1.**
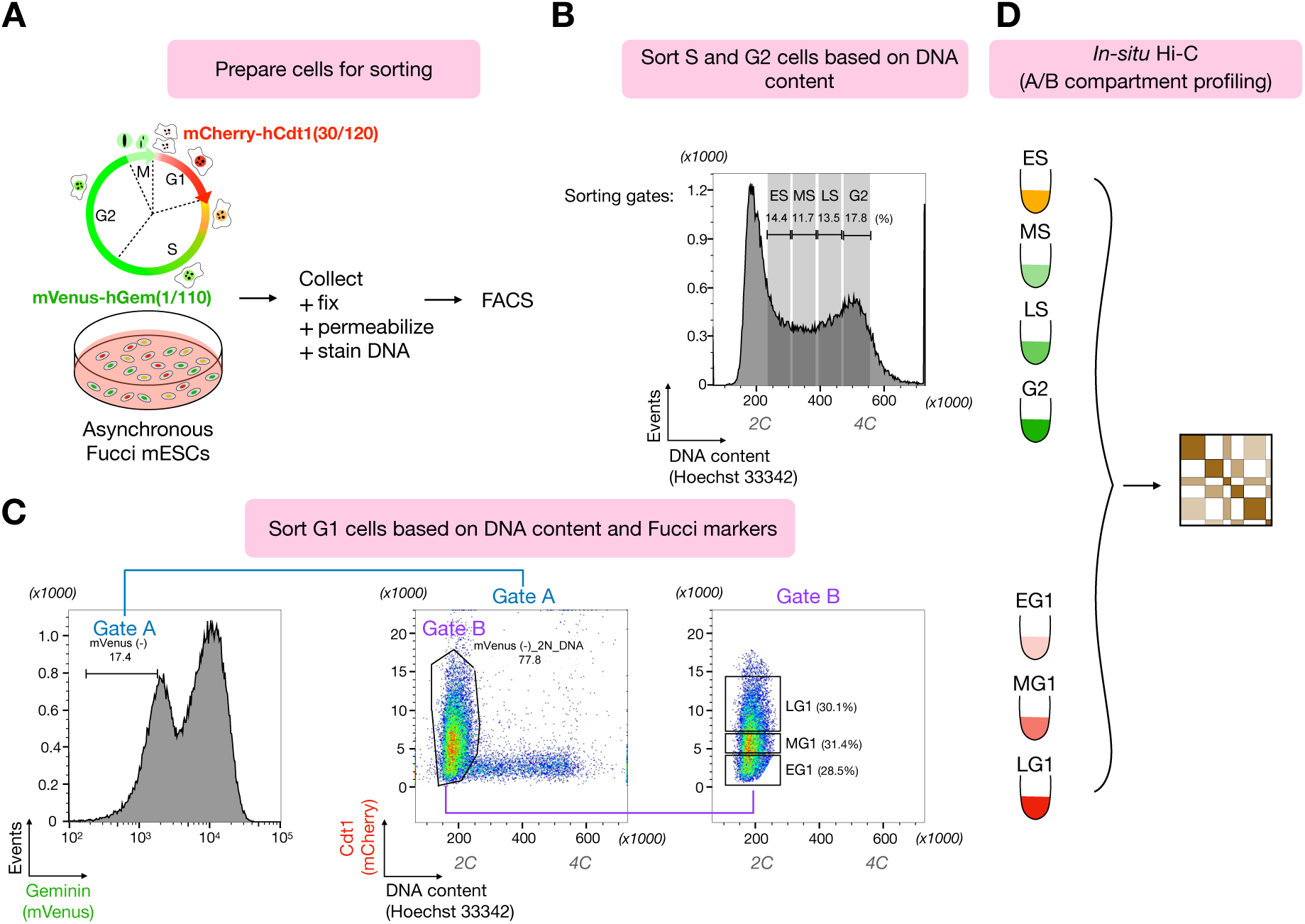
Experimental workflow of cell-cycle phase sorting and Hi-C in mESCs. **(A)** The Fucci2 reporter (top) labels cell cycle phases: G1/early S (mCherry-hCdt1; red) and S/G2/M (mVenus-hGeminin; green). Asynchronous mESC cultures expressing this reporter (bottom) were fixed, permeabilized, and DNA-stained for FACS sorting. **(B)** FACS-sorting strategy for the S and G2 phase cell populations based on DNA content. **(C)** Sequential gating strategy to isolate G1 subpopulations: selection of Geminin-negative cells (Gate A), followed by gating on 2C DNA content (Gate B; entire G1 population), and final fractionation into early, mid, and late G1 based on progressively increasing levels of Cdt1-mCherry fluorescence intensity (approximately 30% of the total population per fraction). **(D)** Sorted cells from each defined phase were subject to in-situ Hi-C. EG1, early G1; MG1, mid G1; LG1, late G1; ES, early S; MS, mid S; LS, late S.

To confirm the suitability of the Fucci markers for this purpose, we performed imaging and fluorescence-activated cell sorting (FACS) analysis following EdU staining, which labels S-phase cells (**Fig. S1A**). As expected, EdU incorporation correlated with Geminin signal accumulation and Cdt1 signal depletion **(Fig. S1B,C)**.

To monitor cell-cycle progression of individual cells, we conducted time-lapse imaging over a 50-hour period, capturing images every 10 minutes using Differential Interference Contrast (DIC), mCherry (Cdt1), and mVenus (Geminin) fluorescence **(Fig. S1D).** We manually tracked individual cells and included only those that completed a full cell cycle from one mitosis to the next for quantitative analysis **(Fig. S1E)**. Our analysis revealed minimal variation in overall cell-cycle length among cells (**Fig. S1F**). We also quantified the duration of the G1 phase for each cell, which we defined as the time from mitosis until mCherry fluorescence reached its peak. The remaining time was defined as the duration of the S/G2/M phase. The G1 phase lasted approximately 1.5 hours (**Fig. S1G**), consistent with the known characteristically short G1 phase of mESCs^18^, with relatively small cell-to-cell variability **(Fig. S1E,G)**.

The above results confirmed that Fucci markers could reliably delineate cell-cycle phases, allowing us to sort homogeneous populations by FACS for subsequent Hi-C analysis. To achieve this, asynchronously growing cells were first fixed, permeabilized, and stained for DNA before being subjected to FACS (**Fig. 1A**). S-phase cells (subdivided into early, mid, and late S, each representing approximately 3-hour time windows, assuming a 10-hour S-phase) and G2-phase cells were sorted based on DNA content (**Fig. 1B**). For G1 cells, we employed a sequential gating strategy using both Fucci markers and DNA content (**Fig. 1C**). The population was first gated to include only Geminin-negative cells, followed by a second gate selecting only cells with a 2C DNA content. From this refined G1 population, we further sorted early, mid, and late G1 fractions based on ascending mCherry fluorescence intensity, with each fraction comprising approximately 30% of the population to achieve sub-hour temporal resolution. Finally, the sorted cells (0.5–1 million cells per fraction) were subjected to in situ Hi-C (**Fig. 1D**).

### Stepwise changes in higher-order chromatin organization during interphase

Hi-C biological replicates were highly concordant (**Fig. S2A,B**). Consistent with previous studies, we observed a continuum of cis-interactions throughout interphase, as shown by contact decay profiles^12^ (**Fig. S2C**). Cis-compartment structure and boundaries (Hi-C PC1) were already distinguishable from early G1 (**Fig. 2A, Fig. S2D**), albeit with a low PC1 contribution rate at this stage (**Fig. S2E**). Furthermore, Hi-C interaction maps (**Fig. 2A, Fig. S3A**) revealed that long-range chromatin interactions formed gradually from early G1 to late G1, expanding away from the diagonal on the map, as previously described in M-to-G1 time-course Hi-C studies^8,9,11^.

**Figure 2.**
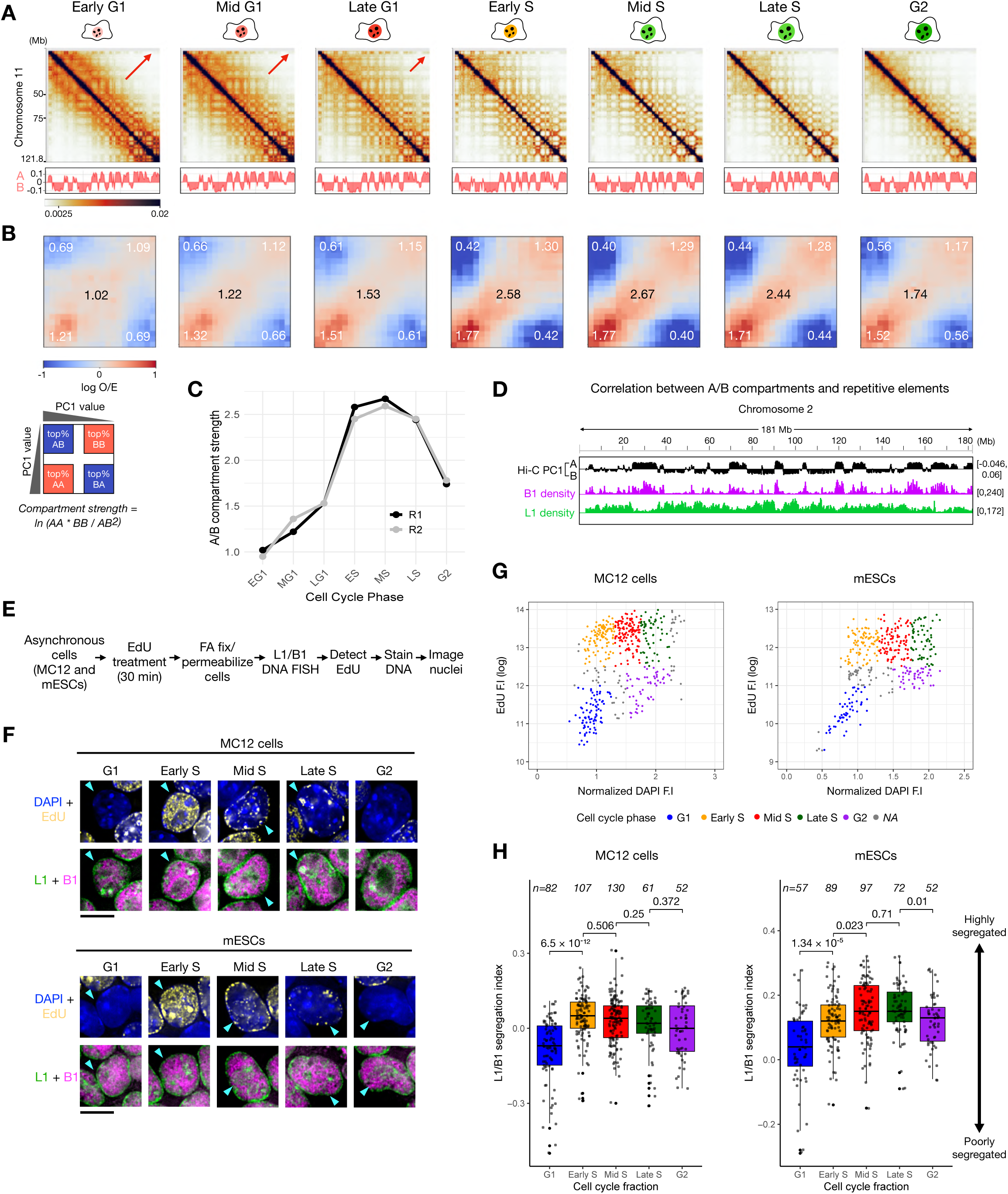
Cell-cycle-phased Hi-C reveals a stepwise progression of nuclear compartmentalization. **(A)** Hi-C contact maps (1-Mb resolution) of chromosome 11 for each cell-cycle phase, with the corresponding A/B compartment profiles (Hi-C PC1) shown below each map. Arrows on the Hi-C maps highlight the progressive outward expansion of contact signal from the diagonal, indicating strengthening of long-range interactions. **(B)** Hi-C saddle plots showing contact enrichment between 1-Mb genomic bins, where both axes are sorted by Hi-C PC1 value (strongest A to strongest B compartment). The schematic below defines the axis ordering. Overall compartment strength (black numerical values) and specific AA, BB, and AB interaction strengths (white numerical values) are quantified. The color scale represents observed/expected (O/E) contact frequencies in 5-percentile increments. Data shown in panels **(A)** and **(B)** are from biological replicate 1 (representative of N=2 biological replicates). **(C)** Hi-C-based compartment strength dynamics across the cell cycle. Each line represents an independent biological replicate. **(D)** IGV browser view illustrating the correlation between A/B compartment profiles (asynchronous mESCs) and L1/B1 repeat densities (mouse chromosome 2, 200-kb resolution). **(E)** Workflow of the L1/B1-EdU FISH experiment (see Methods for details). **(F)** Representative images of MC12 (top) and CBMS1 mESC (bottom) nuclei at the indicated cell-cycle stages, classified based on DAPI and EdU signals. For each nucleus, two merged views are shown: DAPI with EdU, and L1 with B1. Scale bar = 10 µm. **(G)** Single-cell scatter plots of EdU versus DAPI intensity for MC12 cells (left) and CBMS1 mESCs (right). Colors indicate cell-cycle phase assignments as described in the Methods. **(H)** L1/B1 segregation index distributions across cell-cycle stages for MC12 cells (left) and mESCs (right). The segregation index was defined as the negative Pearson’s correlation coefficient between L1 and B1 signals, such that higher values indicate stronger spatial segregation (see Fig. S5B and Methods for details). Data from two biological replicates were merged. n= number of nuclei. Pairwise *P*-values were obtained using a two-tailed unpaired Student’s t-test. EG1, early G1; MG1, mid G1; LG1, late G1; ES, early S; MS, mid S; LS, late S.

Interestingly, however, this expansion from the diagonal culminated in late G1, when interactions spanning the farthest genomic distances (i.e., the regions most distant from the diagonal) were most pronounced (**Fig. 2A, Fig. S3A**). This trend was also evident in the contact decay profiles (**Fig. S2C**) and contact probability plots (**Fig. S2F, Fig. S3B**), which showed the greatest enrichment for contacts > 50 Mb in late G1. As cells entered S-phase, the Hi-C plaid pattern showed a distinct increase in sharpness, which subsequently became progressively blurred during late S-phase and further in G2. Notably, G2 cells displayed a thickened signal along the Hi-C map diagonal (**Fig. 2A, Fig. S3A**), a chromatin organization reminiscent of mitotic chromosomes^5,6^. This finding suggests that the onset of mitotic chromosome condensation may occur earlier than previously thought, during G2 phase (as discussed further below).

While a previous study has linked DNA replication to stronger compartmentalization^12^, we found that the change in compartment strength was strikingly abrupt during the late G1 to early-S transition, as revealed by saddle plots, which quantify Hi-C contact enrichment between genomic bins ranked by their PC1 values (**Fig. 2B,C**). This abrupt late G1-to-early-S shift, which we refer to as ‘compartment maturation’, was accompanied by a notable depletion of A-B interactions and an enrichment of A-A interactions (**Fig. 2B**). Compartment strength then stabilized during S phase, which constitutes the longest phase of the mESC cell cycle (**Fig. S1C**, > 60% cells in S-phase by FACS). Subsequently, a decrease in compartment strength began in late S, followed by an even more significant decrease in G2 (**Fig. 2B,C**). This mirrored the observation from the Hi-C maps mentioned earlier (**Fig. 2A, Fig. S3A**).

The observed decrease in compartment strength during G2 contrasted with a previous report by Nagano *et al.*^12^, which showed maximal compartmentalization in G2. However, it should be noted that a unique compartment identification method was employed in that single-cell Hi-C study^12^, distinct from the standard *de novo* compartment calling approach used in most Hi-C studies (see **Supplementary Note** for details).

Therefore, we attribute this discrepancy not to differences in biological samples or chromatin conformation capture technologies, but rather to technical or interpretative differences arising from the distinct A/B compartment identification methods used. To further support this conclusion, we reanalyzed the cell cycle-phased pooled single-cell Hi-C data from Nagano *et al.*^12^ using the conventional compartment-calling method and observed a trend consistent with our findings (**Fig. S4**), with compartment strength being maximal in early/mid S and decreasing in late S/G2. Thus, our analysis reconciles the apparent discrepancy with Nagano *et al.*^12^ and supports a revised model in which compartment strength peaks in early S-phase and subsequently weakens toward G2, rather than continuously strengthening throughout interphase.

Notably, the pooled single-cell Hi-C data from Nagano *et al.*^12^ showed a clear diagonal thickening in late S/G2 cells (**Fig. S4C**), in agreement with our observations above (**Fig. 2A, Fig. S3A**). Taken together, these data suggest that the structural transition toward mitotic chromosomes may already be initiated in late G2, a view further supported by a live-cell imaging study of interphase-to-mitosis chromosome dynamics showing prophase-like chromosome axis prealignment during G2^19^.

### Stepwise A/B compartment reorganization during interphase is conserved at single-cell resolution

Although our cell cycle-phasing system resolved interphase progression at high temporal resolution, population Hi-C averages chromatin interaction signals across millions of cells and therefore cannot capture potential cell-to-cell variability in compartment segregation. To determine whether the stepwise changes in A/B compartment organization observed during interphase could also be detected in individual cells using a complementary imaging approach.

It is well-established that repetitive elements, particularly transposable elements, show strong associations with A/B compartments: *Alu* and B1 elements are predominantly associated with A compartment regions, whereas L1 elements are enriched in B compartment regions^20^. This relationship is evident when comparing the PC1 profiles of A/B compartments from asynchronous mESCs with the genomic distribution of L1 and B1 density (**Fig. 2D**).

To directly probe this spatial organization, we performed L1/B1-EdU Fluorescence *In Situ* Hybridization (FISH) imaging experiments on both asynchronous CBMS1 mESCs^21^ and MC12 embryonic carcinoma cells^22^ (as briefly described in **Fig. 2E**). As anticipated from their known compartmental associations, the L1 signal localized predominantly at the nuclear periphery, while the B1 signal was enriched in the nuclear interior (**Fig. 2F, S5A**).

For quantitative image analysis, we carefully selected only interphase cells, excluding mitotic cells (including those displaying prophase-like morphology). For each individual cell, we quantified nuclear DAPI and EdU signal intensities as well as the colocalization index (Pearson’s R) between L1 and B1 signals (outlined in **Fig. S5B**). Subsequently, single cells were categorized into specific cell cycle phases (G1, early S, mid S, late S, and G2) based on their DNA content and EdU incorporation patterns (see methods, **Fig. 2G**). Finally, to quantify the extent of L1/B1 segregation, we defined a segregation index based on a previously described method^20^ as the negative value of their colocalization index.

The single-cell distribution of this L1/B1 segregation index within each cell cycle fraction is shown in **Fig. 2H**. While we observed inherent cell-to-cell heterogeneity and overlap in the segregation index across different cell cycle fractions, our imaging results strongly corroborated the Hi-C-based observation of stepwise compartment reorganization during interphase, including the abrupt late G1-to-early-S transition and the subsequent weakening of compartment segregation toward G2. Although our experimental setup did not permit G1 sub-phasing comparable to Hi-C, the broad range in the observed G1 segregation index in both mESCs and MC12 cells is concordant with Hi-C data showing the gradual re-establishment of compartment organization during this period. We also observed a pronounced and statistically significant shift in compartment strength from G1 to early S. Compartment segregation reached its maximum during S-phase and exhibited a notably narrower distribution than in G1 and G2, highlighting reduced cell-to-cell heterogeneity and stabilization of compartment strength. Additionally, the weakening of compartment segregation in G2 was confirmed in our imaging analysis and was not attributable to M-phase cell contamination, as mitotic cells were excluded from the experiment. However, it is possible that the earliest prophase cells were included given that DAPI signal alone is not a foolproof method for their removal; nonetheless, such cells would constitute a minority of the analyzed population. Collectively, these orthogonal findings provide strong evidence that our Hi-C data accurately captures the temporal dynamics of higher-order chromatin organization throughout interphase and that these dynamics are conserved beyond mESCs.

### Compartment maturation requires S-phase entry but is independent of active DNA synthesis

Intrigued by the abrupt and pronounced compartmentalization shift, or compartment maturation, observed during the short transition from late G1 to early S, we investigated its relationship to DNA replication.

Specifically, we asked whether the G1-to-S transition was the direct cause of compartment maturation, or whether the shift in compartmentalization reflects an intrinsic chromatin program that proceeds independently of cell-cycle stage. To test this, we designed two distinct cell-cycle perturbation experiments to induce prolonged arrest at two critical points, either just before or at the start of S-phase, and assessed their effects on compartment architecture.

Our first experimental perturbation aimed to determine whether prolonged G1-arrest, preventing S-phase entry altogether, would influence compartment maturation. Preventing S-phase entry in mESCs is particularly challenging; however, we found that the mTOR inhibitor INK-128 induces a reversible G1-like arrest in mESCs at a concentration of 1 µM^23^ (**Fig. S6A,B,C**). Cells were treated with INK-128 for 24, 72, and 120 hours (**Fig. 3A**). FACS analysis confirmed that cells remained arrested at a 2C DNA content at all timepoints. Concurrently, they exhibited a Geminin-negative state, while maintaining high levels of Cdt1, features characteristic of late G1 cells (**Fig. S6D**). To minimize technical artifacts arising from copy-number differences, cells with a 2C DNA content were sorted at all three timepoints prior to performing Hi-C (**Fig. 3A, Fig. S6D**).

**Figure 3.**
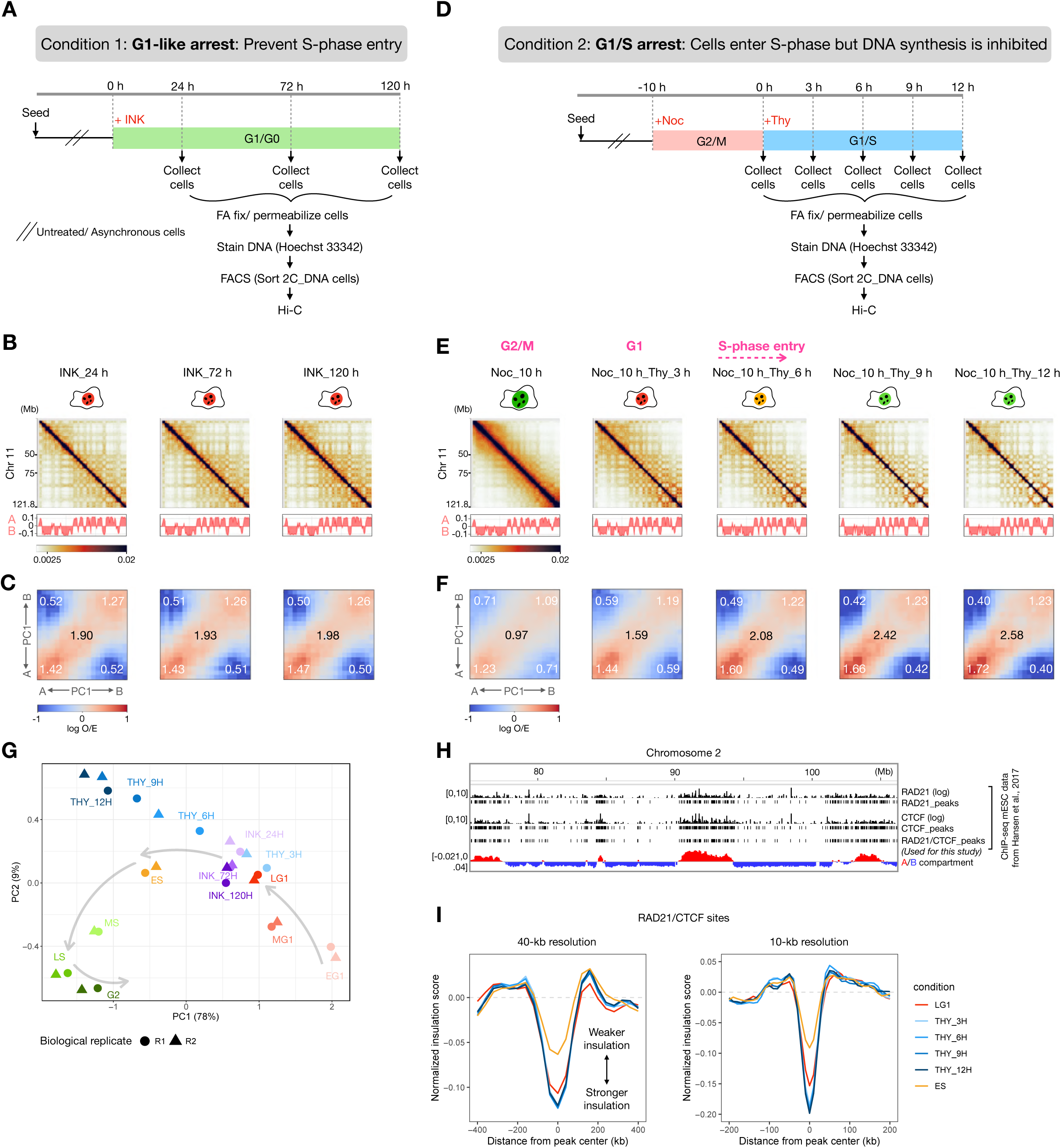
Compartment maturation requires S-phase entry but is independent of active DNA synthesis and cohesin-mediated loop extrusion. **(A)** Experimental design for time-course Hi-C following treatment with INK-128 (INK). **(B)** Hi-C contact maps (1-Mb resolution) of INK-128-treated cells (24, 72, and 120 h), with corresponding A/B compartment profiles (Hi-C PC1) shown below each map. **(C)** Hi-C saddle plot analysis of compartment strength for the data in (B), quantifying the overall strength (numerical values in black) and specific AA, BB, and AB interaction frequencies (numerical values in white). The color scale represents O/E contact frequencies in 5-percentile increments. **(D)** Experimental design for time-course Hi-C following mitotic arrest with Nocodazole (Noc) and release into a Thymidine (Thy) block. **(E)** Hi-C contact maps and A/B compartment profiles for the experiment in (D). Cell-cycle stages (labeled in pink) were inferred for each population based on Fucci2 reporter fluorescence from FACS analysis. **(F)** Hi-C saddle-plot analysis of compartment strength for the data in (E). **(G)** Principal component analysis (PCA) of Hi-C matrices (1-Mb resolution) from asynchronous cell-cycle phases, Nocodazole-Thymidine-blocked cells (THY), and INK-128-treated time-course samples (INK). PCA sample size=50,000. **(H)** IGV browser view of a representative region on chromosome 2 showing ChIP-seq tracks for RAD21 signal, RAD21 peaks, CTCF signal, CTCF peaks, dual-occupancy RAD21/CTCF peaks (ChIP-seq data from Hansen *et al.* 2017^26^), and A/B compartments from asynchronous mESCs (this study; 200 kb resolution). **(I)** Meta-plots showing genome-wide average insulation scores centered at RAD21/CTCF co-occupied sites (1000 randomly sampled peaks) for late G1 (LG1), early S (ES), and G1/S-arrested (THY) cells (at the indicated time points following release from nocodazole and thymidine treatment). Lower insulation scores indicate stronger local insulation and are interpreted as reflecting stronger cohesin-mediated loop extrusion and loop anchoring. Insulation scores were calculated at two resolutions: left, 40- kb resolution using 200- kb sliding windows (±400 kb from peak center); right, 10- kb resolution using 50- kb sliding windows (±200 kb from peak center). Data shown in panels **(B)**, **(C)**, **(E)**, and **(F)** are from biological replicate 1 (representative of N=2 biological replicates). Data in **(I)** is from merged biological replicates (N=2).

Hi-C contact maps from INK-128-treated cells (**Fig. 3B**) resembled a late G1-like chromatin organization (**Fig. 2A**). Saddle plot profiles (**Fig. 3C**) similarly showed compartment strength comparable to that of late G1 (**Fig. 2B**). Although INK-128-treated cells displayed a slightly higher degree of compartment segregation (**Fig. 3C**) and increased long-range interaction formation relative to typical late G1 cells (**Fig. S6E**), the characteristic upward shift in compartment segregation and the enrichment of A-A interactions, indicative of compartment maturation, were notably absent.

The second experiment examined the effect of prolonged G1/S arrest on chromatin compartments. mESCs were first arrested at metaphase using Nocodazole and then released into medium containing a high concentration of Thymidine. Cells were subsequently collected at 3-hour intervals post-release (3, 6, 9, and 12 hours) (**Fig. 3D**). FACS profiles (**Fig. S6F**) confirmed the accumulation of cells with a 2C DNA content following release from Nocodazole arrest. Approximately 6 hours after release, cells began to enter S-phase, as indicated by their transition from a Cdt1-positive/Geminin-negative (Cdt1+Geminin–) to a Cdt1-negative/Geminin-positive (Cdt1–Geminin+) state (**Fig. S6F**). This S-phase fraction progressively increased, leading to an almost complete conversion of 2C DNA content cells to the Cdt1–Geminin+ state after 12 hours of Thymidine treatment, with virtually no remaining G1 (Cdt1+Geminin–) cells. These results confirm that the cell population was successfully synchronized at the G1/S boundary, representing a state where the replication program (including origin licensing, replisome assembly, and helicase activation) has been initiated, as indicated by cell-cycle markers, but ongoing DNA synthesis (elongation) is blocked.

As in the previous time-course experiment, cells collected at all timepoints were FACS-sorted based on their 2C DNA content prior to performing Hi-C **(Fig. 3D, Fig. S6F)**. Hi-C analysis of these samples revealed the emergence of a distinct plaid pattern as early as 3 hours post-release from Nocodazole (**Fig. 3E**), with compartment strength (**Fig. 3F**) comparable to that of late G1 (**Fig. 2A,B**). Strikingly, despite sharing the same 2C DNA content, compartment strength continued to increase over time, as evidenced by the progressive sharpening of the Hi-C plaid pattern (**Fig. 3E**) and by saddle plot analysis (**Fig. 3F**). By 9 and 12 hours of Thymidine treatment, A/B compartment strength had reached levels similar to those of normal early- to mid-S cells (**Fig. 2A,B**). Furthermore, contact probability plots comparing late G1 and Thymidine-treated cells (**Fig. S6G**) revealed an ‘S-like’ trajectory, characterized by a decrease in long-range interactions relative to late G1.

Principal Component Analysis (PCA) of the Thymidine- and INK-128-treated Hi-C matrices, alongside those from normal cell-cycle phases, further clarified these findings (**Fig. 3G**). Specifically, INK-128-treated samples clustered closely with late G1, whereas Thymidine-treated cells aligned closely with the S-phase trajectory, particularly along the PC1 axis, which accounted for 78% of the total variation. Additionally, t-distributed stochastic neighbor embedding (t-SNE) followed by k-means clustering (k=2) of saddle plot data (**Fig. S6H**) revealed a clear separation between INK-128 and Thymidine samples. Thymidine-treated cells (with the exception of the ‘THY_3H’ sample, which largely represented G1 cells) clustered with S-phase samples (thus S-like), whereas INK-128-treated cells grouped with G1 (thus G1-like).

Collectively, our results demonstrate that maturation of nuclear A/B compartments is a direct consequence of S-phase entry, and that this structural reorganization occurs independently of active DNA synthesis.

### Compartment maturation is independent of cohesin-mediated loop extrusion

It has been proposed that cohesin-mediated loop extrusion and A/B compartment strength are often anti-correlated^24,25^. We therefore asked whether compartment maturation could be explained by changes in loop extrusion.

To address this, we first mapped high-confidence cohesin (RAD21) and CTCF co-occupied sites in mESCs using published chromatin immunoprecipitation sequencing (ChIP-seq) data^26^ (**Fig. 3H**). As a functional readout of cohesin-mediated loop extrusion, we quantified insulation at these sites at two resolutions: 40 kb with a 200-kb sliding window to capture TAD boundaries and 10 kb with a 50-kb sliding window to capture fine-scale loops (**Fig. S7A**). Given the high concordance of our cell-cycle-phased Hi-C biological replicates, we merged them for all downstream analyses. Lower insulation scores correspond to stronger local insulation and are generally interpreted as reflecting stronger cohesion-mediated loop extrusion and loop anchoring, whereas higher insulation scores indicate weaker extrusion and less effective anchoring.^27^.

During normal cell-cycle progression, insulation profiles at RAD21/CTCF sites at both resolutions (**Fig. S7B**) showed that insulation scores reached their lowest levels in late G1, as indicated by the steepest meta-plot profile, increased sharply upon entry into S phase, and then gradually declined thereafter. Given that lower insulation scores correspond to stronger local insulation, this pattern indicates that local insulation at RAD21/CTCF sites is strongest in late G1, becomes weakened abruptly in early S phase, and gradually recovers thereafter (consistent with Nagano *et al.*^12^). These dynamics were also reflected in the distribution of insulation scores at these sites (**Fig. S7C**). Thus, under normal cycling conditions, there is indeed an anti-correlation between compartment strength and cohesin-mediated loop extrusion at the G1/S transition: compartment strength increases while insulation decreases.

Importantly, however, this anti-correlation was not systematic. In G1/S-arrested cells (following nocodazole and thymidine block), the decrease in insulation from late G1 to early S was not observed; instead, insulation increased slightly relative to late G1 (**Fig. 3I, Fig. S7D,E**), despite clear compartment maturation at the same time points (**Fig. 3E,F**). Thus, decreases in insulation can be completely uncoupled from increases in compartment strength.

We therefore conclude that the reduction in insulation observed during normal S phase at cohesin/CTCF sites is not a prerequisite for compartment maturation, but is instead likely a consequence of DNA synthesis, potentially reflecting the sister chromatid cohesion function of cohesin. Together, these findings argue against loop extrusion dynamics as the primary driving force underlying compartment maturation.

### Formation of a consolidated A compartment in S-phase

To characterize compartment maturation in greater detail, we performed subcompartment analysis using the Calder2 tool^28^. This computational framework integrates information from multiple principal components of the Hi-C contact matrix to classify genome structures into eight distinct chromatin states based on their multi-dimensional interaction signatures. The resulting subcompartments are ordered by Calder2 rank values ranging from 0.125 (strongest B) to 1 (strongest A), providing a finer-resolution view of the organization within global A/B compartments.

Visual inspection of subcompartment profiles for individual chromosomes at 40-kb resolution (**Fig. 4A, Fig. S8**) revealed that, from early S-phase onward, minor variations among A-subcompartment domains were largely lost and replaced by more homogeneous domains enriched for high-rank, strong A (dark red) states. Quantification of genomic bins assigned to each subcompartment rank revealed a striking and progressive accumulation of rank-1 (strong A) bins from early G1, peaking in mid S-phase and decreasing thereafter in late S and G2 (**Fig. 4B**). Concurrently, medium-ranked (ranks 0.5, 0.625, and 0.75; weak A and B) bins were reduced from early G1 to mid S, suggesting a global shift of subcompartments towards stronger A states (**Fig. 4B**). Consistent with this interpretation, comparison of average subcompartment domain sizes between late G1 and early S phase (**Fig. 4C**) revealed that only high-rank A-subcompartment domains increased significantly in size, with the most statistically significant change observed for rank 0.875 and 1 (strong A) subcompartment domains.

**Figure 4.**
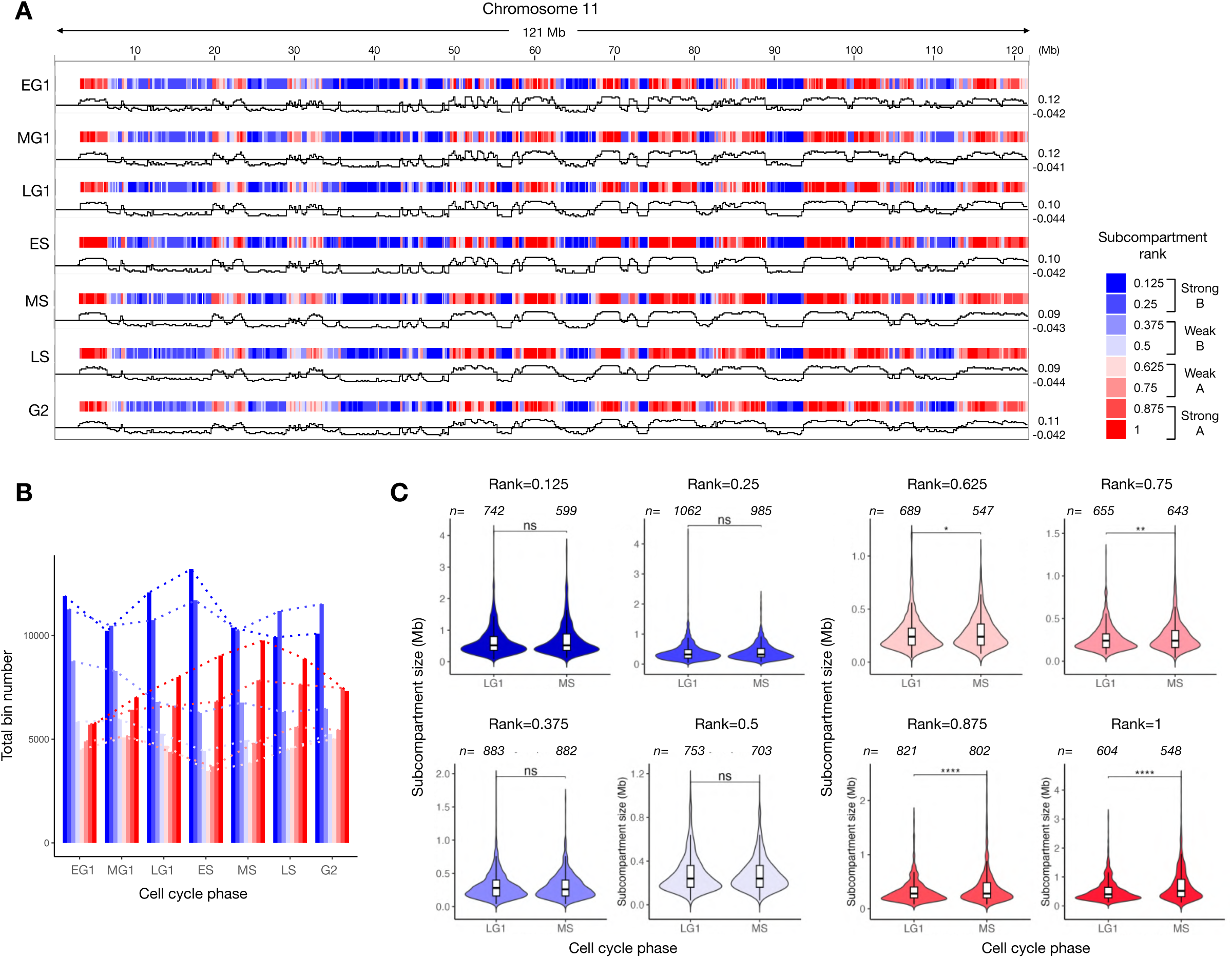
S-phase A-compartment consolidation revealed by subcompartment analysis. **(A)** IGV browser tracks of Calder subcompartments (40-kb resolution) for the entire chromosome 11 across all cell-cycle stages, from early G1 (EG1) to G2. **(B)** Abundance of each Calder subcompartment rank, quantified as the total number of 40-kb genomic bins per cell-cycle stage. **(C)** Violin plots with overlaid box plots showing the size distribution of Calder subcompartment domains in late G1 (LG1) versus mid-S (MS) phase. Domains are defined as contiguous stretches of genomic bins having the same subcompartment rank. Statistical significance was determined using the Wilcoxon rank-sum test; ns, not significant; \**P* ≤ 0.05; \*\**P* ≤ 0.01; \*\*\**P* ≤ 0.001; \*\*\*\**P* ≤ 0.0001. Data in all panels are from merged biological replicates (N=2).

We also observed a “smoother” appearance of A-compartment profiles (Hi-C PC1 > 0) in S-phase cells, with fewer signal spikes (example in **Fig. S9A**). To quantify this trend, we calculated the mean-square gradient (MSG), in which larger values reflect spikier signals and lower values indicate smoother signals (**Fig. S9B**; see Methods). The MSG ratio between strong A versus B compartments showed that, in early G1, the A-compartment signal was more than twice as irregular as the B-compartment signal. This ratio steadily declined after S-phase entry, and by late S and G2, the two signals had converged to a similar degree of smoothness (**Fig. S9C**). Comparison of the average MSG values for the strongest A and B compartment regions (**Fig. S9D**) revealed no consistent trend for the B compartment. In contrast, the A compartment progressively became smoother, reaching its lowest MSG value (smoothest) in mid-S before increasing again in late S and G2.

Together, these results reveal a previously unrecognized feature of chromatin organization during S-phase progression, whereby the A compartment dynamically reorganizes into a more uniform and consolidated strong A-compartment state. We refer to this process as “A-compartment consolidation.”

To test whether this phenomenon is unique to pluripotent cells or more broadly conserved, we analyzed the single-cell HiRES (Hi-C and RNA-seq employed simultaneously) dataset from Liu *et al*.^29^ spanning multiple stages of mouse embryonic development. We selected four cell types with sufficient Hi-C coverage for both G1 and mid-S phases: neural ectoderm (E7.5), neural tube (E8.5), mixed late mesenchyme (E10.5), and extra-embryonic endoderm (E7.5) (**Fig. S10A**; see **Table S1** for read coverage and cell numbers). This selection captured a developmental continuum extending beyond the stage equivalent to mESCs, which is ∼E4.5 epiblast.^30^. Pseudo-bulk Hi-C analysis recapitulated the increase in compartment strength and A-A interactions observed in mESCs (**Fig. S10B**), even in cell types in which B-B interactions were more prominent than A-A interactions in G1. Calder subcompartment analysis at 200-kb resolution (**Fig. S10C**) similarly revealed enrichment of rank-1 (strong A) bins and depletion of medium-ranked (ranks 0.5, 0.625, and 0.75; weak A and B) bins in mid S-phase cells. Together, these findings support the notion that compartment maturation, and the accompanying A-compartment consolidation are not unique to mESCs and may represent a broadly conserved feature of mammalian chromatin organization.

### A-compartment consolidation during S-phase involves enhanced long-range contacts and structural reorganization

To further characterize A-compartment consolidation, we performed distance-resolved compartment strength analysis using the Pentad tool^31^. Specifically, we quantified interactions within A (A-A), within B (B-B), and between A and B (A-B) compartments across four genomic distance ranges throughout interphase: short-range (1–10 Mb), mid-range (10–25 Mb), mid/long-range (25–50 Mb), and long-range (> 50 Mb) (**Fig. 5A**). As expected, positive interaction frequencies were observed for A-A and B-B pairs, and negative frequencies for A-B pairs, at all stages. More notably, we detected a marked enrichment of long-range A-A interactions emerging in early S-phase, which was substantially reduced in G2. Concurrently, short- and mid-range A-B interactions were depleted in early S-phase and remained reduced through mid-S, followed by a modest recovery in late-S and a more pronounced increase in G2. In contrast, B-B interactions exhibited minimal changes across all genomic distances throughout interphase.

**Figure 5.**
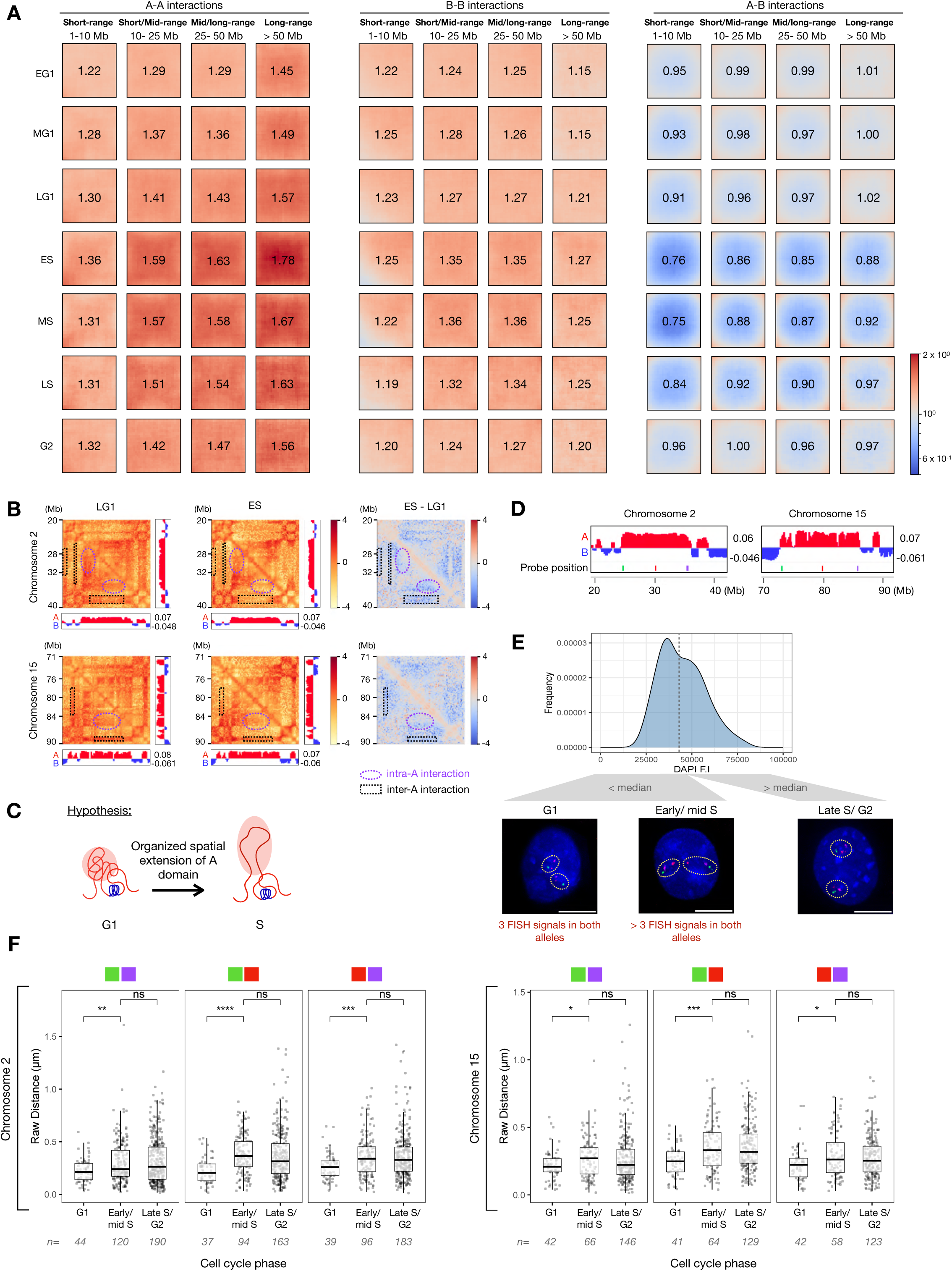
S-phase A-compartment consolidation involves enhanced long-range contacts and structural reorganization. **(A)** Cis-by-distance Pentad plots^31^ for all cell-cycle stages from early G1 (EG1) to G2 phase for short (1–10 Mb), short-mid (10–25 Mb), mid/long (25–50 Mb), and long-range (> 50 Mb) interactions. The value at the center of each plot indicates the mean observed/expected (O/E) contact frequency. **(B)** Observed/Expected Hi-C matrices of two representative regions (chr2: 20–40 Mb and chr8:117–131 Mb) in late G1 (LG1; left panel) and early S phase (ES; middle panel), with corresponding PC1 compartment profiles. The right panel shows a differential heatmap (ES – LG1, early S – late G1). Purple circles highlight weakened interactions between boundaries and the center of a large A domain, or intra-A interactions; black rectangles highlight decreased interactions between neighboring A domains, or inter-A interactions. Panels **(A)** and **(B)** present data from merged biological replicates (N=2) analyzed at 200-kb resolution. **(C)** Schematic model proposing the “A - peninsula” formation, in which internal regions of large A compartments extend away from their boundaries during S-phase. **(D)** Genomic positions of BAC probes targeting selected A domains on mouse chromosomes 2 and 15. Left boundary probes are shown in green, middle probes in red, and right boundary probes in magenta. **(E)** Workflow for assigning cell- cycle stages to nuclei following DNA-FISH. Nuclei were first separated into two groups based on median DAPI signal intensity. Nuclei with above-median DAPI intensity were classified as late S/G2 phase. Nuclei with below-median DAPI intensity were further classified as G1 or early/mid S phase based on the number of FISH spots per allele counted manually (see Methods for details). Representative images for each cell- cycle stage are shown; yellow circles indicate individual alleles. Scale bar = 10 µm. **(F)** Boxplots of pairwise inter-probe distances (µm) by cell- cycle stage. Left panel: chromosome 2; right panel: chromosome 15. For each chromosome, the three probe pairs are (from left to right): green–magenta, green–red, and red–magenta, as indicated above the boxplots. Data from merged biological replicates (N=2); n= nuclei number. Pairwise comparisons between cell cycle stages were performed using the Wilcoxon rank-sum test. P-values were adjusted for multiple comparisons using the Bonferroni method. ns, not significant; *, P ≤ 0.05; **, P ≤ 0.01; ***, P ≤ 0.001; ****, P ≤ 0.0001.

By measuring intra-compartment (intra-A and intra-B; i.e., short-range A-A and B-B) and inter-compartment (inter-A and inter-B; i.e., long-range A-A and B-B) interaction strengths (**Fig. S11A**), we found that all interaction types followed a general “rise-and-fall” trend, similar to that observed in non-distance-resolved saddle plots (**Fig. 2B,C**), with the most pronounced and statistically significant changes occurring between late G1 to early S phase. Most notably, inter-A interaction strength exhibited the greatest and most statistically significant increase during this transition (*P* = 1.74 × 10⁻¹⁰, Δ median = 0.64) (**Fig. S11A,B**), consistent with the enrichment of long-range A-A interactions observed during S-phase (**Fig. 5A**). Together, these observations point to large-scale rearrangements within A-compartment regions as a potential driver of the global shift in A/B compartment segregation at the G1/S transition.

To investigate the structural basis of this phenomenon, we visually inspected Hi-C contact maps of large A-compartment domains in late G1 and early S-phase. This analysis revealed a striking pattern of spatial reorganization within large A domains during S phase, characterized by a sharp reduction in interactions between A-compartment domain boundaries and their internal regions (**Fig. 5B**, purple dotted circles). Most notably, these large A domains appeared to become selectively inaccessible to neighboring smaller A domains, while interactions near domain boundaries were preferentially retained (**Fig. 5B**, black rectangles). The recurrence of this pattern across multiple genomic regions (**Fig. S11C**) suggested a non-random and potentially stereotyped spatial reorganization specific to S phase. We hypothesize that this reorganization process reflects the formation of “peninsula-like” structures, in which internal A-compartment regions extend outward from their boundaries (**Fig. 5C**).

To test this “A peninsula” hypothesis, we performed DNA-FISH in asynchronous mESCs using BAC probes spanning two representative A-compartment domains on chromosomes 2 and 15 (**Fig. 5D**, **Fig. S12A**). Cell-cycle stages (G1, early/mid S, and late S/G2) were assigned to each nucleus based on DNA content and FISH probe numbers, as illustrated in **Fig. 5E**. Pairwise distances between probe centroids were measured per allele in each nucleus (see Methods), and the resulting distance distributions were compared across cell-cycle stages (**Fig. 5F**, **Fig. S12B**).

For both chromosomes, all raw pairwise distances increased significantly from G1 to early/mid S phase, whereas no significant changes were observed from early/mid S to late S/G2. The most pronounced increase was detected between the left boundary probes (green) and the central probes (red), consistent with physical extension of the region containing the red probes.

Together, these DNA-FISH data provide supportive evidence that the spatial reorganization observed in Hi-C maps reflects physical extension of A-compartment domains during S phase. However, we also note substantial cell-to-cell heterogeneity and cannot rule out the possibility that additional factors contribute to the Hi-C interaction pattern change observed during G1 to S-phase transition (**Fig. 5B**, **Fig. S11C**).

### 3D modeling reveals the structural basis of cell-cycle-dependent A/B compartment dynamics

To obtain a holistic structural view of cell-cycle-dependent A/B compartment dynamics, we performed quantitative 3D modeling of chromosomes across all sampled interphase stages using LorDG, a deterministic genome modeler within the GenomeFlow pipeline that generates reproducible polymer-like 3D structures from Hi-C data^32^. Analysis of merged biological replicates at 200-kb resolution using optimized parameters (conversion factor = 0.6, max iterations = 10,000) yielded models that closely matched the experimental Hi-C data, with an average absolute Spearman correlation of 0.77 (**Fig. S13**).

Visually, the simulations with overlaid A/B compartment tracks at the same resolution (e.g., chromosomes 2 and 17 in **Fig. 6A**) faithfully recapitulated the transition from rod-like early G1 chromosomes to a more interphase-like organization in late G1. Most importantly, the model captured the emergence of extended “peninsula-like” structures within the A compartment regions during early S phase. In G2, the models showed a more uniform folding pattern accompanied by overall chromosomal compaction (**Fig. 6A**).

**Figure 6.**
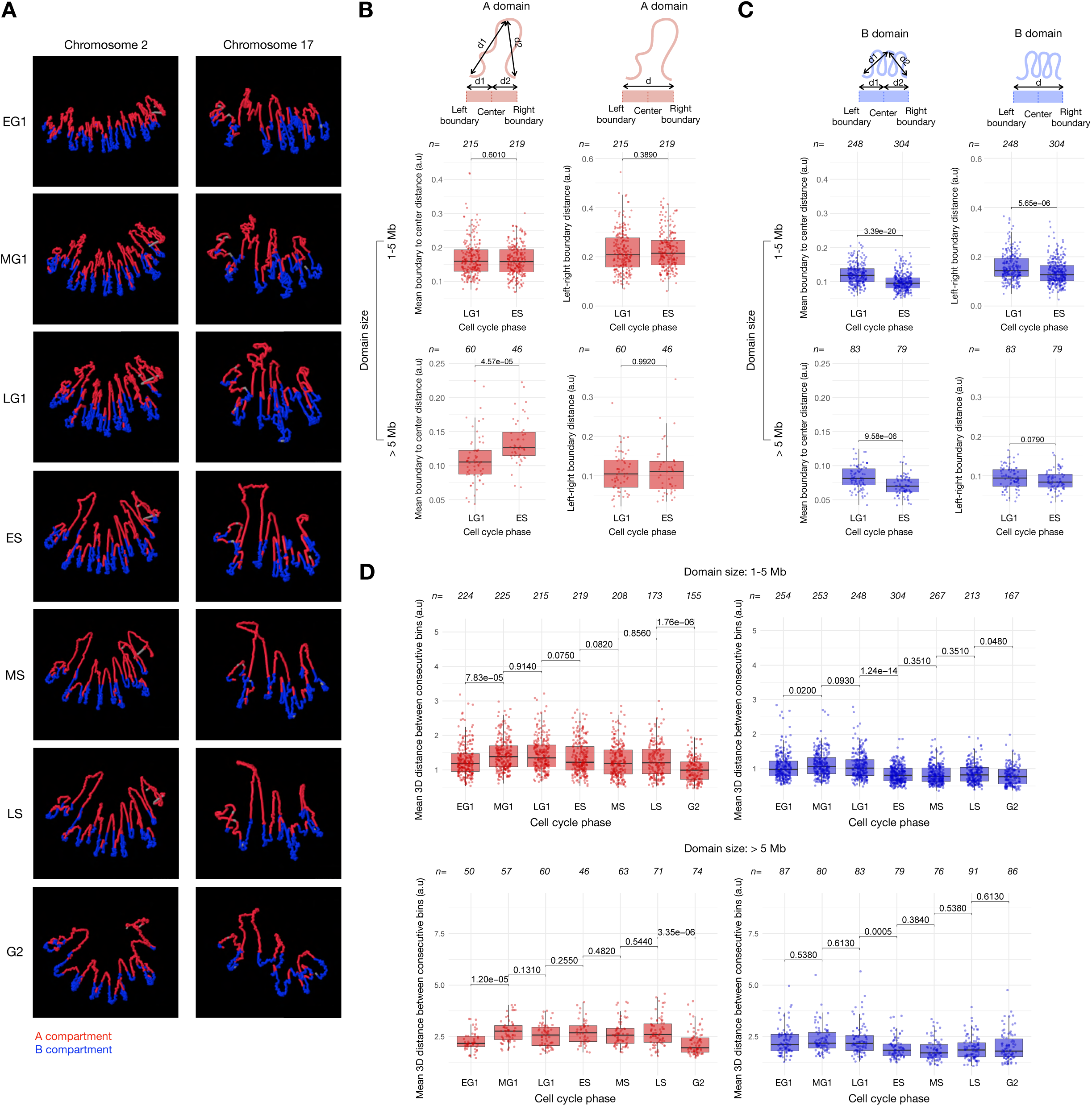
3D genome modeling from Hi-C recapitulates temporal interphase compartment dynamics. **(A)** 3D genome structures of chromosomes 2 and 17 across the cell cycle, simulated from Hi-C data using the LorDG Modeler in GenomeFlow^32^ using a conversion factor of 0.6 (10,000 iterations). **(B,C**) (Left panels) Quantification of outward extension for compartments A and B, showing the mean shortest Euclidean distance from the domain center to its boundaries (calculated as (d1 + d2)/2), normalized by the number of bins per domain. (Right panels) Quantification of boundary movement, showing the shortest Euclidean distance between the boundaries of adjacent domains, normalized by the number of bins per domain. Analyses are shown for small (1–5 Mb) and large (> 5 Mb) domains, comparing late G1 and early S phases. **(D)** Distribution of the mean bin-to-bin distance within A (Left panels) and B (Right panels) compartment domains across cell-cycle phases. Data are shown for small (1–5 Mb) and large (> 5 Mb) domains. Boxplots in **(B–D)** show the median and quartiles. Each point in the scatter represents a single domain (n = total domains). Pairwise comparisons between consecutive phases were calculated using the Wilcoxon rank-sum test, and p-values were adjusted using the Benjamini-Hochberg method. Data are from merged biological replicates (N=2), analyzed at 200-kb resolution.

To quantitatively assess the formation of these “peninsula-like” structures in a manner analogous to our DNA-FISH analysis (**Fig. 5F**), we measured global A-domain extension, defined as the size-normalized shortest domain boundary-to-center distance. Analysis of small (1–5 Mb) and large (> 5 Mb) domains revealed that only large A domains showed a significant increase in global extension from late G1 to early S-phase (**Fig. 6B, left panels**). Importantly, this extension occurred without changes in the inter-boundary distance (**Fig. 6B, right panels**), supporting a model in which internal chromatin regions extend outward from their boundaries from late G1 to early S, a global spatial reorganization specific to domains larger than 5 Mb.

We next asked whether B compartment domains underwent similar global structural changes. In stark contrast to A domains, B domains exhibited significant decreases in both boundary-to-center and boundary-to-boundary distances across all domain sizes (1–5 Mb and > 5 Mb), consistent with global chromatin compaction during the late G1-to-early S transition (**Fig. 6C**).

Having established that both A and B domains undergo substantial global reorganization during the late G1-to-early S transition, we next examined whether these structural changes in A and B domains were accompanied by alterations in local chromatin compaction (i.e., changes in the physical packing of the polymer-like chromatin fiber). To assess this, we calculated the average bin-to-bin distance within size-stratified domains as a proxy for local compaction levels. Decreases or increases in this metric reflect tighter or looser local chromatin packing, respectively, whereas stable values would indicate unchanged local packing at this scale.

In A-compartment domains, significant changes in local compaction levels were detected only between early and mid G1 (decompaction) and between late S and G2 (compaction), with compaction levels converging to a uniform state across A domains in G2 (as indicated by the minimal dynamic range), reaching levels comparable to those of B domains (**Fig. 6D, Fig. S14A,B**). Notably, however, A-domain local compaction remained essentially unchanged from late G1 through S phase despite substantial global structural reorganization of these domains. Thus, the emergence of “peninsula-like” structures in large A domains reflects global positional rearrangements rather than changes in local chromatin compaction.

As expected, B domains were generally more compact than A domains across all cell-cycle stages (**Fig. 6D, Fig. S14A**). However, their most significant local compaction change occurred specifically between late G1 and early S, becoming more uniformly compact at the onset of S-phase (**Fig. 6D, right panels; Fig. S14C**). These findings imply a previously unrecognized active contribution of B-compartment domain reorganization in S-phase-dependent compartment maturation.

Taken together, these results refine our understanding of A/B compartment structural dynamics throughout interphase. The simulations not only recapitulated known features, such as chromosome decompaction from early to late G1, but also uncovered previously underappreciated aspects of compartment maturation during S-phase. Specifically, they revealed global changes in the spatial trajectory of A-domain chromatin occurring without accompanying changes in local chromatin compaction, providing a structural explanation for the gain of long-range A-A interactions and the emergence of consolidated A-compartment domains observed during S-phase (described in the previous section). Moreover, our analyses revealed that B domains are also actively engaged in compartment maturation, showing more uniform compaction locally and globally in early S, a feature previously overlooked by conventional Hi-C analyses. Finally, the models indicate that G2 phase is characterized by relatively uniform local chromatin compaction across the genome. We note that these models are derived from population-averaged Hi-C data and should therefore be interpreted as a heuristic framework for understanding A/B compartment dynamics, rather than as definitive representations of individual nuclei.

## Discussion

In this study, we established a simple cell-sorting and chromatin-profiling strategy that precisely resolves cell-cycle phases across the entire interphase. Using this system, we captured cell populations with exceptionally high temporal resolution, achieving sub-hour resolution within the G1 phase (**Fig. 1, Fig. S1**). By applying comprehensive A/B compartment-centered analyses to these carefully staged populations, we obtained a nuanced view of how nuclear organization is dynamically remodeled throughout interphase. This approach allowed us to propose a revised model of cell-cycle-dependent chromatin dynamics that challenges the prevailing view (**Fig. 7**). Our model delineates a stepwise cell-cycle-phase-dependent reorganization of nuclear compartmentalization, comprising four distinct stages. The first stage, chromosome unfolding, occurs during progression from early to late G1 phase as chromatin expands from its highly compacted mitotic state. This is followed by compartment maturation, characterized by an abrupt enhancement of A/B compartment separation upon entry into S-phase. The third stage, compartment stabilization, persists throughout S-phase, whereas the final stage, chromosome refolding, occurs during G2 as cells prepare for mitosis. Our primary focus was the previously overlooked maturation stage. We found that this reorganization is a direct consequence of S-phase entry but occurs independently of ongoing DNA synthesis. A defining feature of this stage is the emergence of a consolidated A compartment, involving extensive spatial reorganization of individual A domains and the establishment of extensive long-range A-A interactions.

**Figure 7.**
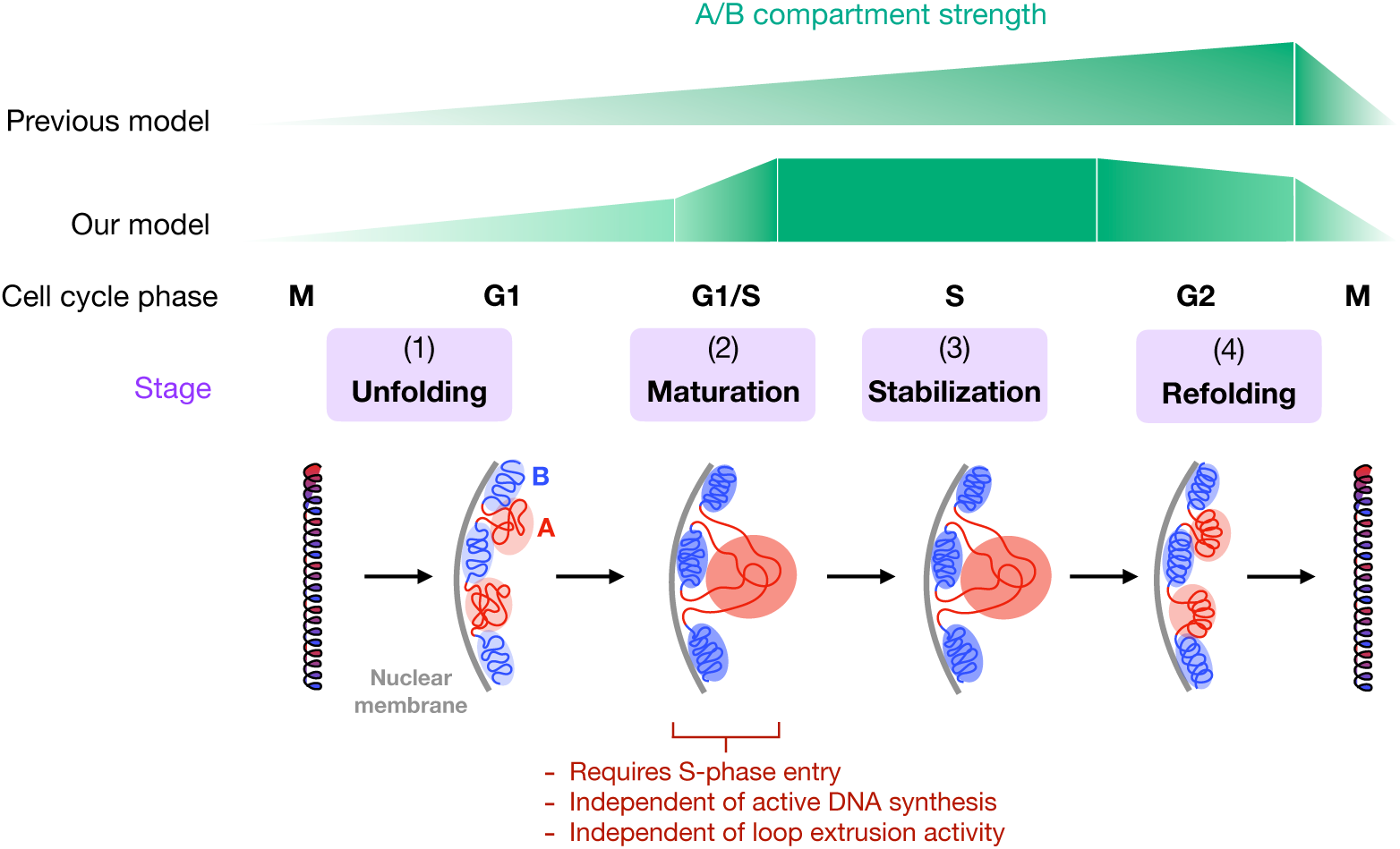
Model of stepwise 3D genome reorganization during the cell cycle. The model proposes four sequential stages: (1) Chromosome unfolding (G1): Gradual formation of long-range interactions from the compact mitotic state. (2) Compartment maturation (G1/S Transition): An abrupt enhancement of compartmentalization upon S-phase entry, independent of DNA synthesis and cohesin-mediated loop extrusion. This stage is characterized by A-compartment consolidation, accompanied by increased long-range A-A interactions and uniform compaction of the B compartment. (3) Compartment stabilization (S-phase): The matured compartment state is maintained throughout the S phase. (4) Chromosome refolding (G2): Global compaction begins in preparation for mitosis.

Our findings provide several key conceptual insights. While the M-to-G1 and G2-to-M phase transitions are well known for their dramatic chromatin reorganization, we contend that the G1-to-S transition is equally critical yet remains significantly underexplored. This transition may therefore represent a particularly informative time window for discovering and characterizing novel 3D genome regulators. More broadly, our results identify S-phase entry as a major driving force of large-scale 3D genome reorganization, prompting reconsideration of how replication and chromatin architecture are functionally linked.

The prevailing model posits that replication timing is a readout of a pre-established nuclear framework defined by relatively stable compartments and domains^14,33–35^. However, our data indicate that this hierarchy warrants re-evaluation. We propose a model of dynamic interplay, in which chromatin changes associated with the G1-to-S transition actively reshape the very architectural landscape from which the replication program itself is derived. Our results also reveal an unexpected degree of plasticity in A/B compartment organization even within a single cell type, a feature that may be underappreciated due to the predominant focus on Hi-C PC1 of asynchronously growing cell populations when assessing A/B compartments. More generally, these findings underscore that, while the field increasingly pursues ever-higher data resolution at sub-TAD and loop levels, important principles of genome organization remain to be uncovered at larger spatial scales. High-resolution approaches, though valuable, should not overshadow the importance of coarse-grained perspectives of chromatin organization, as demonstrated by our findings.

At the same time, recent ultra-high-resolution Hi-C studies^36,37^ have revealed compartment-like features at much finer genomic scales, emphasizing that A/B compartmentalization is, to some extent, inherently scale-dependent. Understanding how these fine-scale, often transient micro-compartments relate to the more stable, large-scale segregation patterns described here will be an important direction for future studies. In addition, it is important to note that our Hi-C analyses, although based on highly resolved and precisely staged cell populations, remain ensemble measurements. Future studies combining ultra-high temporal resolution with single-cell analysis will be essential to resolve the precise kinetics and cell-to-cell heterogeneity of these cell cycle-dependent chromatin reorganization events.

The biological significance of these dynamic S-phase chromatin behaviors remains an important open question. One possibility is that A-compartment reorganization facilitates DNA replication. A-compartment regions are gene-rich, euchromatic, and replicate early in S-phase. Their dynamic reorganization during early S phase may therefore enhance the accessibility or spatial coordination of replication origins and associated factors. It is tempting to speculate that this reorganization process may support the formation of coupled replication factories, in which sister replisomes function as a single coordinated unit^38^, potentially facilitating efficient replication initiation and smooth progression of replication forks. Such structural plasticity may help optimize replication efficiency and ensure faithful execution of the replication timing program.

A second possibility is that this remodeling improves coordination between replication and transcription. Given that A-compartment regions are also transcriptionally active, their reorganization during replication may help alleviate topological conflicts between transcriptional and replication machineries, allowing both processes to proceed efficiently. Moreover, rearrangement of A domains could effectively expand their interface with nuclear bodies such as splicing speckles, thereby enhancing spatial compartmentalization in ways that support efficient DNA replication. These interpretations are broadly consistent with recent work demonstrating that the relative positioning of genomic regions with respect to nuclear locales influences genome organization, replication timing regulation, and gene expression^39^.

Uncovering the biological significance of these chromatin dynamics first requires identifying the molecular mechanisms responsible for compartment maturation in S-phase. Components of the replication machinery, particularly those not directly involved in DNA synthesis itself, represent especially intriguing candidates.

Their potential to influence higher-order chromatin organization is supported by recent findings, including evidence that certain replisome components in *Drosophila* undergo phase separation^40^ and that replisome loading can restrict chromatin motion independently of DNA synthesis^41^. Beyond the core replisome, other S-phase-associated factors and molecular processes such as transcriptional machinery, chromatin modifiers, architectural proteins, and nuclear organizers are also plausible candidates^42,43^. Elucidating the precise roles and interplay of these factors will be an important next step in understanding the mechanistic basis of compartment maturation.

While our work primarily focused on the critical G1-to-S transition, events occurring during the S-to-G2 transition warrant equal attention. We speculate that mitotic chromosome reformation may begin in G2, potentially earlier than currently appreciated, thereby facilitating efficient chromosome segregation. Such early initiation may minimize disruption to ongoing nuclear processes. Accordingly, the S/G2 boundary represents another promising window for investigating cell-cycle-dependent chromatin reorganization.

In summary, our findings support a dynamic and cell-cycle-aware model of A/B compartmentalization, in which chromatin architecture is continuously remodeled and precisely tuned to accommodate and coordinate essential nuclear functions at appropriate stages of the cell cycle.

## Materials and methods

### Cell culture

The R26p-Fucci2 mESC line was obtained from the RIKEN BDR animal facility, LARGE (derived from the R26p-Fucci2 mouse line, Acc. No. CDB0203T). To perform accurate flow cytometry (FACS) color compensation for this dual-reporter line, we also derived control mESC lines from single-reporter mice. For this, oocytes from R26-mCherry-hCdt1(30/120) (Fucci-red, Acc. No. CDB0264K) and R26-mVenus-hGeminin(1/110) (Fucci-green, Acc. No. CDB0265K) mice were fertilized and cultured *in vitro* at the LARGE facility to the morula/blastocyst stage. The zona pellucida was removed using Tyrode’s solution (Sigma, T1788). Individual blastocysts were plated onto a feeder layer of mitotically inactivated mouse embryonic fibroblasts (MEFs) in ES cell derivation medium. The medium was composed of KnockOut^TM^ DMEM (Thermo Fisher Scientific, 10829018) supplemented with 20% knockout serum replacement (KSR, Gibco,10828-028), 1x MEM Non-Essential Amino Acids (Nacalai, 06344-56), 1x EmbryoMax® Nucleosides (Millipore, ES-008-D), 0.5 mg/mL penicillin-streptomycin (Nacalai, 09367-34), 100 µM beta-mercaptoethanol (Gibco, 21985-023), 1 µM PD0325901 (Wako, 162-25291), 3 µM CHIR99021 (Wako, 034-23103), and 1000 U/mL leukemia inhibitory factor (LIF, Nacalai, NU0012-2). After 5–7 days of incubation at 37°C with 5% CO₂, the inner cell mass outgrowth was dissociated and replated. Emerging mESC colonies were selectively picked, expanded, and passaged to establish stable cell lines.

The mouse ESC lines R26-mCherry-hCdt1 (Fucci-red), R26-mVenus-hGeminin (Fucci-green), R26p-Fucci2, and CBMS1 (non-fluorescent) were maintained in 2i/LIF medium. This medium consisted of NDiff 227 base medium (Cellartis, Y40002) supplemented with the following: the two inhibitors (2i) PD0325901 at 1 μM and CHIR99021 at 3 μM; 0.5 mg/mL penicillin-streptomycin; 0.1 mM 2-mercaptoethanol; and LIF at 1000 U/mL. Cells were cultured on iMatrix-511 (Matrixome, 892012)-coated dishes at 37°C with 5% CO₂ and passaged every 2–3 days using trypsin/EDTA (Nacalai, 32777-44).

The MC12 cell line^22^ was maintained in high-glucose DMEM (Thermo Fisher Scientific, 11965092) supplemented with 10% fetal bovine serum (FBS). Cells were cultured at 37°C in a humidified atmosphere with 5% CO₂ and passaged every 2–3 days.

### Flow cytometry

Cell sorting and analysis were conducted using a Sony MA900 cell sorter. The following laser and filter configurations were used: mVenus (488-nm laser, 525/50 nm filter), mCherry (561-nm laser, 617/30 nm filter), Hoechst 33342 and FxCycle Violet (405-nm laser, 450/50 nm filter), and Alexa Fluor 647 (638-nm laser, 665/30 nm filter). For color compensation, formaldehyde-fixed CBMS1 mESCs served as a double-negative control. Single-color controls were established using formaldehyde-fixed Fucci-red mESCs, Fucci-green mESCs, and, where applicable, EdU(AF 647)-labeled CBMS1 mESCs. Data were analyzed using FlowJo (v10.8.1; BD Life Sciences).

### Live-cell imaging of Fucci mESCs

Fucci mESCs were plated at a density of 0.1 million cells/mL in a 12-well plate and allowed to adhere for several hours prior to imaging. Live-cell imaging was performed for 50 hours using the Cell<u>D</u>iscoverer 7 automated microscope (Zeiss), acquiring images every 10 minutes. Imaging was conducted with a 20x objective (NA 0.5) using the system’s automated focus maintenance. Fluorescence of the Fucci reporters was captured using the following settings: mCherry was excited with a 590 nm LED and emission was collected with a 635/50 nm bandpass filter; mVenus was excited with a 520 nm LED and emission was collected with a 540/25 nm bandpass filter. Simultaneous transmitted light images were acquired using both DIC and Phase Contrast modalities. For the analysis of the live-cell imaging data, manual tracking of nuclei was performed using the TrackMate plugin (v6.0.30)^44^ in Fiji (ImageJ2, v2.16.0)^45^ using DIC and phase contrast images (tracker radius was set to 5 pixels). Only cells that completed a full cell cycle, defined as starting two frames after cell division and ending at the last frame before the subsequent cell division, were analyzed. The mean fluorescence intensities (mVenus and mCherry) for each tracked nucleus were exported using the TrackMate Extras plugin. Subsequent data processing and analysis were performed in R (v4.3.2; R Core Team, 2025).

Briefly, the data from two biological replicates were merged, and cell cycle trajectories were aligned to the start of G1 phase. Fluorescence signals were baseline-corrected by subtracting the minimum intensity for each channel per cell. This pipeline was used to quantify cell cycle length, G1 length (timing of when cells reach maximum mCherry intensity), and to plot the normalized Fucci fluorescence dynamics.

### EdU labeling and detection

To correlate Fucci markers with S-phase progression, DNA synthesis was assessed using the Click-iT Plus EdU imaging kit (Invitrogen). For imaging, cells were plated on sterile glass coverslips 24 hours prior to experimentation. Cells were pulsed with 10 µM EdU for 15 minutes, then fixed with 4% formaldehyde and permeabilized with 0.5% Triton X-100. The Click-iT reaction was performed according to the manufacturer’s protocol, incubating cells with the Alexa Fluor 647 azide dye for 30 minutes in the dark. Nuclei were counterstained with Hoechst 33342 (1 µg/mL), and coverslips were mounted with Vectashield antifade mounting medium (Vector Laboratories no. H1000). Imaging was performed on a Nikon Ti-E Eclipse microscope equipped with a SOLA SE LED light source and standard filter sets for DAPI, FITC, TRITC, and Cy5. Images were acquired with a 40x objective and analyzed with the Fiji software. For parallel analysis by flow cytometry, cells were harvested and processed in suspension, undergoing the same EdU labeling, fixation, permeabilization, and Click-iT reaction steps using the analogous Click-iT Plus EdU flow cytometry kit (Invitrogen), before being stained for DNA content with 1 µM FxCycle Violet stain.

### L1/B1-EdU FISH

We performed DNA FISH for L1 and B1 genomic repeats coupled with EdU detection, building on a previously described protocol^46^. Briefly, MC12 cells and CBMS1 mESCs were pulsed with 10 µM EdU for 10 min, fixed with 4% formaldehyde in PBS for 10 min at room temperature (RT), washed with PBS, and permeabilized with 0.5% Triton X-100 in PBS for 10 min. After three PBS washes, samples were treated with 0.1 M HCl for 5 min, followed by incubation with 0.1 mg/mL RNase A in PBS for 45 min at 37°C. Cells were then equilibrated in 2× SSC and pre-hybridized in 50% formamide/2× SSCT (2× SSC, 0.1% Tween-20) for 5 min at RT and 20 min at 47°C. Genomic DNA was denatured by floating the dish in an 83°C water bath for 3 min in 50% formamide/2× SSCT. The denaturation solution was immediately replaced with 100 µL of hybridization buffer (2× SSC, 50% formamide, 20% dextran sulfate) containing 0.5 µM each of L1- and B1-specific fluorescently labeled DNA FISH probes (custom synthesis, Thermo Fisher Scientific).

Probe sequences were as follows:

L1-mouse-1_Alexa488: AGGACACATGCTCCACTATGTTCATAGCAG
L1-mouse-2_Alexa488: AGATGCCCATGAACATACAAGAAGCCTACAGAACT
B1-mouse-1_Alexa594: gcctggtctacagagtgagttccaggacag
B1-mouse-2_Alexa594: cagcacttgggaggcagaggcaggcggatt

Hybridization was carried out under a coverslip in a humidified chamber at 37°C for 16 h. Post-hybridization washes included two 15-min washes in 2× SSCT at 50°C, followed by two 1-h washes in 2× SSCT at RT. For EdU detection, cells were incubated with Click-iT reaction cocktail (Click-iT EdU Alexa Fluor 647 Imaging Kit) for 30 min at RT, washed twice with PBST, and counterstained with DAPI (1:300 in PBS) for 15 min. Coverslips were mounted with Vectashield antifade mounting medium. Images were acquired on a DeltaVision Olympus IX71 microscope with a 60× oil immersion objective, with deconvolution and maximum-intensity projection performed automatically using the standard SoftWoRx acquisition software (v.6.5.2).

Nuclei were segmented automatically on the DAPI channel using Fiji, manually curated to exclude edge and mitotic cells, and saved as ROIs. For each nucleus, integrated DAPI and EdU intensities were measured. For L1 and B1 channels, background was subtracted using a rolling ball radius of 30 pixels, followed by default auto-thresholding to generate binary masks. Colocalization between L1 and B1 was quantified within each nuclear ROI using Pearson’s correlation coefficient (Costes regression, 10 randomizations) with the Fiji Coloc 2 plugin. Using R, datasets from replicate experiments were merged and normalized so that the G1 DAPI peak equaled 1. Nuclei with normalized DAPI values outside 0.5–2.5 were excluded. EdU positive and negative populations were defined using log transformed EdU intensity thresholds specific to each cell line: 12.5 for mC12 cells and 11.5 for mESCs. S phase cells were identified as the EdU positive population and subdivided into three equal-width bins by DNA content (for both cell types). G2 cells were defined as the EdU negative population with DAPI between 1.5 and 2.5 (for both cell types). G1 cells were defined as the EdU negative population with DAPI between 0.5 and 1.25. To exclude ambiguous cells near the EdU positivity threshold, a stricter EdU upper bound was applied: 12 for mC12 cells and 11 for mESCs (compared to the EdU negative cutoffs of 12.5 and 11.5, respectively). The L1/B1 segregation index was defined as the negative value of Pearson’s R. Statistical comparisons between consecutive cell cycle phases were performed using two-tailed unpaired Student’s t tests.

### Multicolor DNA FISH for A compartment consolidation analysis

BAC clones were obtained from Hirose Chemicals. For chromosome 2: left boundary (Green, RP23-455B24, chr2:24666611-24852492), middle (Red, RP23-315H12, chr2:30052581-30244450), right boundary (Magenta, RP23-98E3, chr2:35480102-35693633). For chromosome 15: left boundary (Green, RP23-56M12, chr15:73020495-73240968), middle (Red, RP23-458K15, chr15:79799699-79970129), right boundary (Magenta, RP23-409A19, chr15:85714311-85920895). BAC DNA was extracted using alkaline lysis miniprep. Probes were labeled with fluorescence-dUTP (Green-dUTP, Enzo Life Sciences 02N32-050; Red-dUTP, Enzo Life Sciences 02N34-050; Cyanine5-dUTP, Perkin Elmer NEL579001EA) by nick translation (Abbott Molecular 07J00-001). DNA-FISH was performed as described^47^. Briefly, asynchronously growing mESCs or cells treated with 0.01 μg/ml Colcemid (Wako 045-16963) for 2 h (for mitotic chromosome enrichment) were harvested, incubated in 75 mM KCl for 15 min, and fixed in methanol:acetic acid (3:1). Labeled probes, mouse Cot-1 (Thermo Fisher Scientific 18440-016), and salmon sperm DNA (Thermo Fisher Scientific 15632-011) were precipitated, resuspended in hybridization buffer (10% dextran sulfate, 2× SSC, 1% Tween-20, 50% formamide), and denatured at 80°C for 10 min. Fixed cells were dropped onto slides, washed with 2× SSC, treated with RNase A (10 µg/mL in 2× SSC) for 1 h at 37°C, dehydrated through sequential 5-minute washes with 70%, 90%, and 100% ethanol at room temperature before being air-dried at 58°C for 1 hour, then denatured in 70% formamide/2× SSC at 80°C for 3 min. Hybridization was performed overnight at 37°C. Post-hybridization washes were performed in 50% formamide/2× SSC at 45°C (three times) and 0.1% SSC at 60°C (three times). Slides were counterstained with DAPI (500 ng/mL) and mounted with Vectashield. Images were acquired on a DeltaVision Olympus IX71 microscope with a 60× oil immersion objective. Deconvolution and maximum-intensity projection were performed using SoftWoRx (v.6.5.2).

Using Fiji, nuclei were segmented automatically on the DAPI channel using Otsu dark thresholding, binary masking, close gaps, fill holes, watershed separation, and particle analysis (size 10–Infinity). ROIs at borders or with poor segmentation were manually removed. DNA FISH spots were detected using a custom script: after Gaussian blur (sigma = 4), thresholds were applied per channel (Green: 830–65535, Red: 1150–65535, Magenta: 154–65535), and particles between 0.50 and 10.00 pixels were detected and added as ROIs with channel-specific prefixes (“FISH_Green_”, “FISH_Red_”, “FISH_Magenta_”). Undetected spots were added manually. FISH ROIs were assigned to parent nuclei by centroid containment. For each nucleus, a k-means clustering algorithm initialized with the two most distant points defined two allele territories; spots were assigned to the nearest allele with a constraint of maximum one spot per color per allele. Using R, nuclei were separated by DNA content using the median DAPI intensity. Nuclei above the median were assigned as late S/G2. Nuclei below the median were manually assigned to G1, S, or NA based on FISH signal count per allele. For each allele, pairwise distances between FISH signals (Green-Red, Magenta-Red, and Green-Magenta) were calculated. Statistical comparisons between consecutive cell cycle phases (G1 vs S, S vs late S/G2) were performed using two-tailed Wilcoxon rank-sum tests.

### Cell cycle synchronization and collection for Hi-C

For Hi-C experiments, Fucci mESCs were collected under three primary conditions. First, to obtain cells at various cell cycle stages (EG1, MG1, LG1, ES, MS, LS, and G2), an asynchronous population was harvested directly (50-100 million cells per experiment). Second, for the G1/S time-course experiment, cells were synchronized at the G2/M border using a 10-hour treatment with 50 ng/mL Nocodazole (Sigma, M1404). One dish of these synchronized cells was harvested as the G2/M time point. In parallel, other synchronized dishes were washed twice with PBS and released into a medium containing 2.5 mM thymidine (Sigma, T1895); these were then harvested every 3 hours from 3 to 12 hours post-release (collecting 2-4 million cells per time point). Finally, for a G1 arrest time-course, asynchronous mESCs from separate dishes were treated with 1 µM INK-128 (MedChemExpress, HY-13328) and harvested at 24, 72, and 120-hour time points (collecting 2–4 million cells per time point). All collected samples from these procedures were processed for Hi-C and subsequent FACS analysis as described in the following section.

### Sample preparation for Hi-C

All harvested cells were processed for Hi-C using an established protocol^48^ with minor modifications. Briefly, harvested cells were fixed in 1% formaldehyde in PBS for 10 minutes at room temperature, using 1 mL of fixative per 1 million cells. The reaction was quenched with 0.125 M glycine. After permeabilization with 0.1% saponin, cells were stained with Hoechst 33342 (10 µg/mL) for 30 minutes in the dark to label DNA for cell cycle analysis. Cells were then FACS-sorted using a Sony MA900 cell sorter in purity mode. Laser configurations were as previously described in the ‘Flow Cytometry’ section, and the gating strategies for isolating specific cell cycle populations are detailed in the Results section and corresponding figures. Sorted cell pellets were immediately flash-frozen in liquid nitrogen and stored at –80°C until Hi-C and library preparation. Then, Hi-C libraries were prepared from 0.5–1 million fixed cells (frozen pellets) according to an established protocol^48^. In brief, nuclei were isolated, chromatin was digested with DpnII (NEB, R0543), and restriction fragment overhangs were filled with biotinylated nucleotides prior to blunt-end ligation. After reverse crosslinking and DNA purification, 0.5–1 µg of DNA was sheared to 150–400 bp fragments using a Covaris S220 sonicator, with size selection performed using AMPure XP beads (Beckman Coulter, A63881).

Biotin-labeled fragments were captured on Streptavidin-coated magnetic beads (Dynabeads M-280, Thermo Fisher Scientific 11205D), and the resulting libraries were prepared for sequencing using the NGS LTP Library Preparation Kit (KAPA, KK8232). Final libraries were sequenced on a NovaSeq X platform to generate 150 bp paired-end reads (approximately 100 million reads per sample; range 70–130 million).

### Generation of genome-wide L1/B1 repeat density tracks for IGV visualization

To create genome-wide density tracks for L1 and B1/Alu elements in the mm9 reference genome, RepeatMasker annotations were downloaded from the RepeatMasker^49^ repository. We generated 200 kb genomic windows using bedtools^50^ makewindows. L1 and B1/Alu repeats were counted per window using bedtools intersect -c, and the resulting counts were converted to .bedGraph format.

### Hi-C data processing and quality control

Hi-C sequencing reads were processed using the ‘FANC’ pipeline^51^ (v0.95) (https://github.com/vaquerizaslab/fanc) with the command: ‘fanc auto <fastq1> <fastq2> <output> -g mm9 -i <bwa_index> -r MboI -n -b 1mb,200kb -t 30 -f --fanc-parallel -tmp -q 30’. This command performs alignment to the mm9 genome using BWA (v0.7.17), enables iterative mapping (--iterative) with a quality cutoff of 30 (-q 30), and produces a valid pairs file (.pairs) of aligned reads alongside unbinned .hic files. These were then used to generate normalized (-n) contact matrices at 1-Mb and 200-kb resolutions (-b 1mb,200kb). When applicable, biological replicates were merged into a single contact map using ‘fanc hic <file1> <file2> <output> -b 1mb -a -n --restore-coverage’, where the -a flag aggregates contacts and --restore-coverage maintains library size after merging. Library quality was assessed using ‘fanc pairs -s’ for mapping statistics and ‘fanc hic -s’ for cis-chromosomal contacts (see **Table S2** for all quality metrics).

### single-cell Hi-C data processing

Multiplexed FASTQ files from Nagano *et al*.^12^ were downloaded from GEO (GSE94489). This dataset consists of pooled single-cell libraries, where cells were sorted and sequenced collectively by cell cycle phase. Data were processed identically to our in-house data, with the exception that iterative mapping was omitted and the restriction enzyme was specified as MboI. A summary of downloaded datasets and processing statistics is provided in **Table S3**.

### HiRES data processing

Public single-cell sequencing data from the Liu *et al*. HiRES study^29^ were obtained from GEO (GSE223917). This dataset contained a mixture of DNA and RNA sequences. Metadata (science.adg3797_table_s2.xlsx) were filtered to select specific cell types and developmental stages (ExE endoderm E7.5, Mixed late mesenchyme E10.5, Neural ectoderm E7.5, Neural tube E8.5) with sufficient sequencing depth (> 100 million reads across G1 and mid S cell populations). Individual cell IDs were extracted, and corresponding FASTQ files were downloaded. Adapter trimming was performed with Cutadapt (v1.18)^52^ using the RNA-specific sequence (GGTTGAGGTAGTATTGCGCAATG) to discard non-ligated RNA fragments (--discard-trimmed) and isolate DNA-derived Hi-C pairs. Read quality was assessed using FastQC (v0.11.9). Per-cell-type replicates were generated by merging forward (_1_DNA.fastq.gz) and reverse (_2_DNA.fastq.gz) reads from individual cells. The resulting merged FASTQ files were once again assessed using FastQC and then processed through the FANC pipeline using the MboI restriction enzyme site, without iterative mapping (see **Table S1** for sample details and statistics).

### Probability (P(s)) plots derivation

Contact probability (P(s)) curves were generated using FANC’s expected value calculation (fanc expected). The analysis was performed on 1-Mb resolution Hi-C matrices, comparing the decay of contact probability as a function of genomic distance (s) for each tested condition. The resulting P(s) curves were plotted to visualize differences in interaction frequency decay across conditions.

### Contact map comparisons

Pairwise comparisons between Hi-C matrices were performed using the ‘fanc compare’ function. Matrices at 200-kb resolution were directly compared using the -Z (z-score normalization) and -I (insulation correction) parameters. The resulting difference matrices were visualized using the ‘fancplot’ function to highlight regions of increased or decreased contact frequency between two conditions.

### Principal component analysis (PCA) of Hi-C matrices

Principal component analysis was performed on 1-Mb resolution Hi-C contact matrices from all biological replicates using FANC’s PCA implementation (fanc pca)^51^, analyzing the top 50,000 genomic bins (default parameter). The analysis included all chromosomes with a minimum distance cutoff of 1 Mb.

### A/B Compartment analysis and saddle plot generation by FANC

A/B compartments were called from Hi-C contact matrices at 1-Mb resolution using the ‘fanc compartments’ command. A Pearson correlation matrix was calculated from the .hic file. The first eigenvector (PC1) was computed from this matrix and its sign was oriented using the GC content of the mm9 genome (-g option). The oriented PC1 and the resulting A/B compartment domain coordinates were output to BED files (-v and -d options, respectively).

Saddle plots were generated from the compartment eigenvector (1-Mb resolution) using the ‘fanc compartments’ command. The -e option was used to output the enrichment plot, and the --compartment-strength option was used to calculate a quantitative compartment strength score (as defined by Flyamer *et al*.^53^ as the natural logarithm of AA * BB / AB^2^). Compartments were binned into 5-percentile intervals using the -p option, and the resulting enrichment matrix was saved to a file with the -m option for downstream analysis.

### t-SNE and K-means clustering of saddle plots

Saddle plot enrichment matrices at 1-Mb resolution were analyzed in R using t-distributed stochastic neighbor embedding (t-SNE) and K-means clustering. Individual matrices were flattened into vectors and dimensionality reduction was performed with perplexity 7 and 500 iterations. K-means clustering (k=2) was then applied to the resulting coordinates to identify groups of samples with similar compartment interaction patterns.

### Format conversion for subsequent Hi-C analysis

Hi-C pairs were converted from FANC’s proprietary format to the standardized .pairs format and processed as individual replicates into multi-resolution cooler matrices (.mcool) using cooler^54^ (v0.9.3) ‘cload pairs’ and ‘zoomify’. When applicable, biological replicates were merged at 10-kb resolution using ‘cooler merge’, and the resulting combined matrix was processed with ‘cooler zoomify’ using identical parameters (resolutions from 10 kb to 1 Mb with iterative balancing) to generate the final merged .mcool file.

### Calculation of contact probability decay profiles

Contact decay profiles were generated from the same standardized .pairs files. A 10% downsampling of each file was performed using ‘pairtools sample’ (v0.3.0)^55^ with a random seed of 1 to ensure reproducibility and facilitate computational efficiency. The resulting downsampled pairs files were processed with a custom R script to calculate the contact probability as a function of genomic distance, adapted from the method described by Nagano *et al.*^12^. For each read pair, the linear genomic separation was calculated for intra-chromosomal (cis) interactions. These distances were log2-transformed and binned. The contact frequency for each bin was normalized by the total number of valid cis reads to generate a relative contact probability. Finally, the decay profiles were visualized as a heatmap using the ‘fields’ package in R, where the x-axis represents different cell cycle phases, the y-axis represents genomic distance, and the color intensity corresponds to the percentage of total contacts.

### Compartment calling at 200-kb resolution

Although compartment calling is available in the standard ‘FANC’ pipeline, we found that compartments were not robustly called at 200-kb resolution. To address this, we performed compartment analysis using a custom Python implementation that executes principal component analysis (PCA) via cooltools (0.3.2)^56^.

Compartments were called from balanced .mcool files using a modified approach that calculates principal components through correlation and covariance matrices before PCA. The first principal component (PC1) was extracted as the compartment track, where positive values correspond to active (A) compartments and negative values to inactive (B) compartments. PC1 sign was oriented using a gene density track for mm9. The analysis calculated three key outputs for each sample: compartment eigenvectors, eigenvalues, and the eigenvalue contribution rate for PC1, which quantifies the proportion of variance in the contact matrix explained by A/B compartmentalization (PC1).

### Hierarchical clustering of compartment profiles

A/B compartment profiles at 200-kb resolution were analyzed using DeepTools (v3.5.5)^57^. First, compartment eigenvectors were summarized using ‘multiBigwigSummary bins’ across all samples. Pearson correlation coefficients were then calculated and visualized as a clustered heatmap using ‘plotCorrelation’ with outlier removal.

### Insulation score analysis at CTCF/RAD21 co-bound sites

Published CTCF and RAD51 ChIP-seq peak files from Hansen *et al.* (2017)^26^ (GSM2418860 and GSM2418859) were downloaded from the European Nucleotide Archive, converted from .txt to BED format, and lifted over from mm10 to mm9 using the UCSC liftOver tool^58^. Overlapping peaks were identified with ‘bedtools intersect’ and deduplicated to generate a final set of high-confidence CTCF/RAD21 co-bound sites. Genome-wide signal tracks (bigWig files) were similarly converted to mm9-compatible bedGraph files for visualisation.

Insulation scores were computed from balanced Hi-C contact maps for each cell-cycle phase using cooltools ‘diamond-insulation’ at two resolutions: 10 kb with a 50 kb sliding window, and 40 kb with a 200 kb sliding window (ignoring the two diagonal bins). Output BED files were cleaned to remove invalid entries, sorted, and converted to bedGraph format.

All subsequent insulation profile (meta-plot) and statistical analyses were performed using R. For meta-plot generation, insulation scores were extracted from a random subsample of 1,000 peaks (set.seed(123)). For each peak, a region spanning ±400 kb (40-kb data) or ±200 kb (10-kb data) around the peak centre was interpolated using linear approximation. Mean insulation scores and standard errors were calculated at each position, and normalised meta-plots were generated by subtracting the average flanking insulation score. For boxplot quantification, insulation scores were extracted directly at the bin center for all high-confidence CTCF/RAD21 sites. Pairwise comparisons between consecutive cell-cycle stages (EG1 vs MG1, MG1 vs LG1, LG1 vs ES, ES vs MS, MS vs LS, LS vs G2) were performed using the Wilcoxon rank-sum test, with p-values adjusted using the Benjamini-Hochberg method.

Multi track genome browser views were generated with pyGenomeTracks^59^ using a configuration file that included Hi-C contact maps, CTCF/RAD21 binding sites, insulation scores, and A/B compartment eigenvectors, visualized consistently across conditions at chr2:63,000,000–77,000,000.

### Quantification of compartment smoothness using mean-square-gradient (MSG)

To quantitatively assess the smoothness of A/B compartment profiles, we calculated the mean-square-gradient (MSG) of the compartment signal (PC1) derived from 200-kb binned data. The MSG is defined as MSG = ⟨(∂f/∂x)²⟩, where f(x) is the compartment signal as a function of genomic position x. This metric quantifies the average rate of change of the compartment signal, where higher values indicate more variable, less smooth signals, while lower values reflect more uniform, smoother compartment signals. The genomic gradient of PC1 was computed for all chromosomes in each cell cycle stage using the ‘numpy.gradient’ function in Python^60^. To focus on core compartment domains while avoiding boundary regions, we selected genomic bins where the compartment signal exceeded one standard deviation above the chromosome mean for A compartments, or fell below one standard deviation for B compartments. The MSG was calculated separately for these strong A (MSG_A,1σ) and B (MSG_B,1σ) regions. To measure the relative changes in compartment signal smoothness throughout the cell cycle and normalize for technical variations, we computed the ratio MSG_A,1σ/MSG_B,1σ.

### Distance-based compartment strength quantification

We used ‘Pentad’^31^ (<u>github.com/magnitov/pentads</u>, commit 66c19f4) to measure A/B compartment interactions at various genomic distances. Hi-C contact matrices in .mcool format and compartment signals (PC1) at 200 kb resolution were used as input for Pentad’s cis-interaction scripts (get_pentad_cis.py) with rescaling to 50 matrix (pixel) bins. Compartment strength was quantified (quant_strength_cis.py), and distance-dependent analyses were performed at 1-10 Mb, 10-25 Mb, 25-50 Mb, and > 50 Mb distances (get_pentad_distance.py). Subsequent analysis in R compared chromosome-specific strength values for inter- and intra-compartment interactions (A-A, B-B, A-B) across cell cycle phases. Delta median values were calculated for consecutive cell cycle transitions to identify the magnitude of change in compartment strength.

### Chromatin subcompartment analysis

Chromatin subcompartments were identified using ‘CALDER2’^28^ (https://github.com/CSOgroup/CALDER2) executed within a Docker container (lucananni93/calder2:latest). For compatibility with CALDER, contact matrices in .mcool format were converted to .hic files using ‘cooler dump’ to export balanced interaction scores, followed by conversion with juicer_tools (v1.22.01)^61^ using the mm9 genome assembly. Analyses were performed at 40-kb resolution for newly generated data and 200 kb for reanalyzed HiRES data, with binning chosen to ensure a strong correlation of compartment rank with a reference dataset (Pearson’s rho > 0.4).

Subcompartment calls were filtered in R to include only genomic bins consistently detected across all cell cycle fractions. Subcompartment sizes were calculated as contiguous genomic regions sharing identical compartment ranks within the hierarchical system spanning eight ranks (0.125–1), representing the continuum from B-compartments (0.125–0.5) to A-compartments (0.625–1).

### 3D Genome structure modeling from Hi-C data

Chromosome structures were reconstructed from Hi-C data using ‘GenomeFlow’ (v2.0) (https://github.com/jianlin-cheng/GenomeFlow) with the ‘LorDG-3D Modeler’ function^32^. We generated intra-chromosomal contact maps from 200-kb resolution Hi-C matrices using ‘fanc dump --only-intra’ and separated them into individual chromosome datasets. For each chromosome, 10,000 iterations of the modeling algorithm were performed with an optimized conversion factor of 0.6 to maximize the correlation between reconstructed 3D distances and original Hi-C interaction frequencies. A/B compartment annotations were derived from Calder subcompartment calls (200-kb resolution) and stitched into continuous A or B domains using custom R scripts. Spatial coordinates (x, y, z) were extracted for each 200-kb genomic bin, annotated with A/B compartment identity, and filtered to retain only bins common to all samples. From these structural models, we quantified 3D compartment organization for domains > 1 Mb using three spatial metrics calculated from the relative 3D Euclidean distances (in arbitrary units, a.u.) in the reconstructed models:

1. Local compaction: The mean 3D Euclidean distance between all consecutive bins within a domain.
2. Domain shape: The average 3D distance from the domain boundaries to its central bin.
3. Inter-boundary distance: The 3D distance between the leftmost and rightmost boundary bins. Boundary-to-center and inter-boundary distances were normalized by the number of bins per domain. Statistical significance was assessed using Wilcoxon rank-sum tests with Benjamini-Hochberg correction.

### Statistics and reproducibility

All statistical analyses are detailed in the figure legends and main text. This includes the specific tests used, the number of independent biological replicates (N), the number of technical measurements or data points (n), and either exact P-values (*P*) or standard significance indicators (ns, not significant; *P* ≤ 0.05; \**P* ≤ 0.01; \*\**P* ≤ 0.001; \*\*\**P* ≤ 0.0001). In box plots, the central line denotes the median, the box boundaries represent the 25th and 75th percentiles, and the whiskers extend to the most extreme data points within 1.5 times the interquartile range from the box.

## Acknowledgements

We are grateful to all members of the Hiratani laboratory for their insightful discussions and feedback throughout this project. We extend special thanks to Y. Furuta, H. Kiyonari, and M. Kaneko (RIKEN BDR animal facility, LARGE) for providing the Fucci2 mESCs and T. Abe for single Fucci reporter blastocysts. We also thank A. Ueno for technical assistance in deriving mESCs from blastocysts, R. Poonperm for experimental and analytical support, Y.K for technical support with L1/B1-EdU FISH experiments, and M. Takiguchi for technical assistance with live-cell imaging on the CellDiscoverer7 microscope. Finally, we thank T. Takahashi, Y. Kawasoe, A. Sakaue-Sawano, A. Miyawaki, Y. Wang, J. Cheng, and M. Magnitov for their valuable scientific input. We also acknowledge the use of large language models, specifically ChatGPT (OpenAI), Gemini (Google), DeepSeek (DeepSeek-AI), and Perplexity AI, to assist with the editing of the manuscript text, as well as for troubleshooting code during data analysis. The authors reviewed, edited, and take full responsibility for all content. This work was supported by a RIKEN International Program Associate (IPA) fellowship to L.C.; RIKEN BDR intramural grants, the RIKEN Pioneering Project ‘Genome Building from TADs’, MEXT KAKENHI Grant Number JP18H05530, JSPS KAKENHI Grant Numbers JP20K20582 and JP25H00982, and JST CREST Grant Number JPMJCR20S5 to I.H.

## Author contributions

L.C. and I.H. conceived the study. L.C. designed and performed the majority of the experimental work and conducted the bioinformatic analyses. Hi-C experiments were performed jointly by L.C. and T.I. H.M. generated key preliminary data and facilitated data conversion between analytical platforms. A.O. characterized the INK-128-induced G1-like arrest in detail and provided experimental support. S.T. designed and performed L1/B1-EdU FISH experiments as well as provided technical assistance for mESC derivation from blastocysts, and R.C. performed MSG analysis and generated the panels in Fig. S9. The manuscript was written by L.C. and I.H. with input from all co-authors.

## Data availability

The raw Hi-C sequencing data have been deposited in the DNA Data Bank of Japan (DDBJ) under BioProject accession PRJDB37948. The processed Hi-C data are available in the Gene Expression Omnibus under accession GSE312348. The custom scripts for Hi-C analyses are available at Zenodo: https://doi.org/10.5281/zenodo.20552820. The live-cell imaging, single-cell tracking data, and analysis script (Fucci-mESCs) are available at Zenodo: https://doi.org/10.5281/zenodo.17509930. The L1/B1-EdU DNA FISH data (Unprocessed images, ROI coordinates, raw measurements, scripts) are available at Zenodo: https://doi.org/10.5281/zenodo.20473330. The A-compartment consolidation DNA FISH data (Unprocessed images, ROI coordinates, raw measurements, scripts) are available at Zenodo: https://doi.org/10.5281/zenodo.20552838. Source data for figures are provided.

## Supplementary materials

### Supplementary note: A comparison of methodologies for A/B compartment analysis

Our study characterizes compartment dynamics using *de novo* identification of A/B compartments from population Hi-C data. This approach differs fundamentally from the single-cell compartment “strength” metric pioneered by Nagano *et al.* (2017)^12^, and the choice of methodology is directly linked to the specific biological question being addressed.

The method developed by Nagano *et al.* was designed to quantify cell-to-cell variation by first defining a stable reference map of A and B domains derived from pooled single-cell data. The compartment strength for each individual cell is then calculated based on how closely its intra-chromosomal contact pattern conforms to this pre-defined reference. This provides a robust measure of structural heterogeneity but inherently assesses fidelity to the population-average architecture, rather than the strength of compartmentalization per se.

In contrast, our *de novo* approach re-identifies compartment identities independently for each cell-cycle-staged population, allowing us to capture the temporal progression of compartment organization, including large-scale rearrangements and the emergence of new organizational states that would otherwise be masked by comparison to a static reference. Thus, while the method of Nagano *et al.*^12^ provides an excellent measure of compartment pattern conservation across cells, our approach is specifically tailored to reveal how these patterns evolve over time during cell-cycle progression.

### Supplementary figure legends

**Figure S1.**
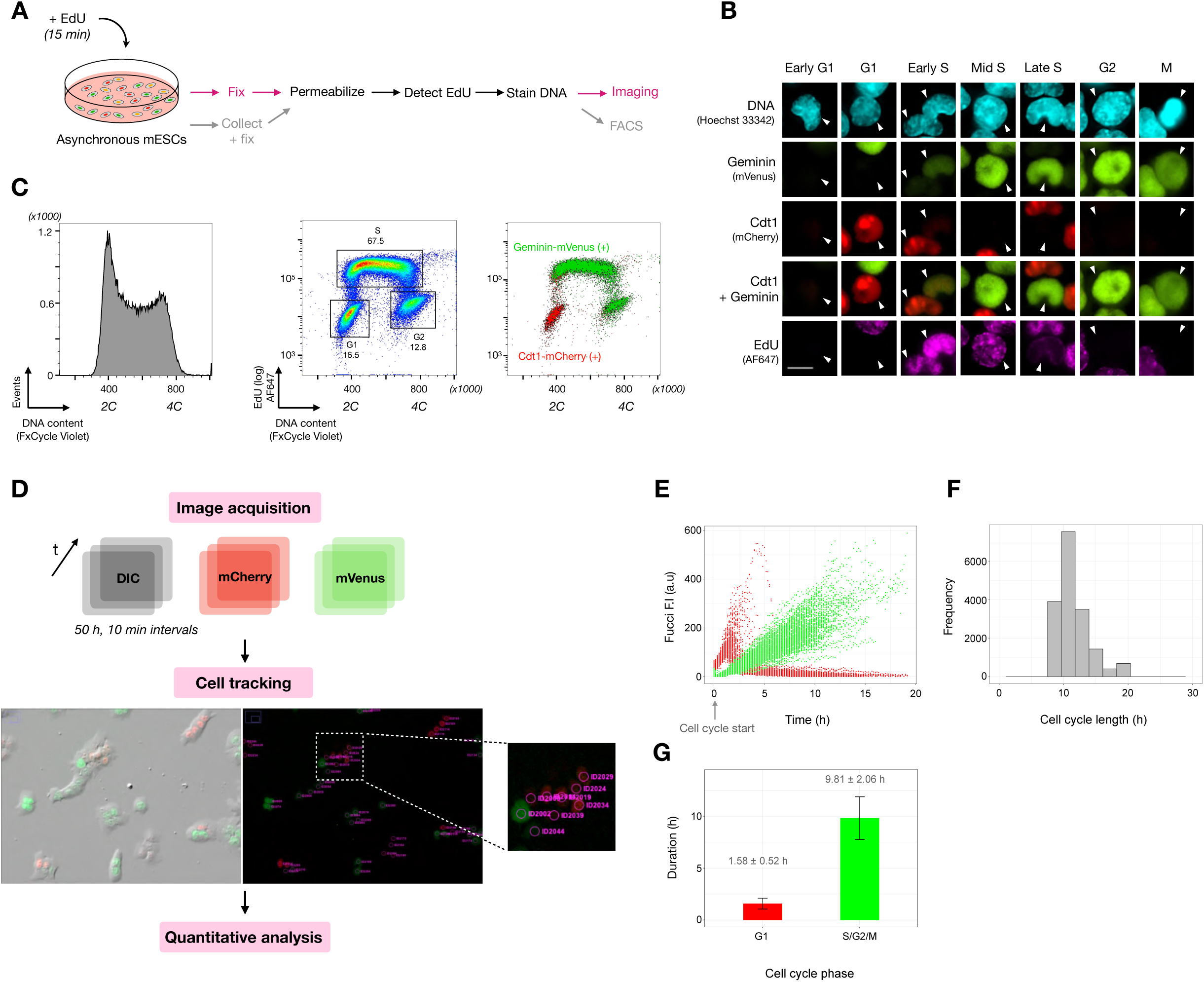
Validation of Fucci cell-cycle reporters by imaging, FACS, and time-lapse analysis. **(A)** Schematic of the EdU labeling protocol in Fucci mESCs for subsequent FACS or imaging analysis. Steps common to both procedures are shown in black. **(B)** Representative images of Fucci-expressing nuclei (indicated by white arrows) after EdU treatment and DNA labeling across G1, S, G2, and M phases. Scale bar = 10 µm. **(C)** FACS analysis of EdU-labeled Fucci mESCs, showing DNA content (left), EdU versus DNA content with gated cell-cycle populations (middle), and the same EdU versus DNA content plot with overlaid Fucci signals (right). Geminin-mVenus (green) accumulates at the G1/S transition, while Cdt1-mCherry (red) is enriched in G1. **(D)** Schematic of the time-lapse imaging and cell-tracking strategy. **(E**) Temporal dynamics of Fucci reporters (Geminin-mVenus, green; mCdt1-mCherry, red) in single cells after baseline correction and alignment to cell-cycle start (two frames post cell division). Mitotic cells are excluded. **(F)** Quantification of total cell-cycle length. The mean duration was estimated to be 10.83 ± 2.49 h. **(G)** Distribution of time spent in G1 (red) versus S/G2/M (green). G1 length was defined as the interval from mitosis to the peak of mCherry fluorescence, with the remaining cell cycle duration assigned to S/G2/M phases. Data in **(E–G)** are from tracked cells across 2 biological replicates (n=131).

**Figure S2.**
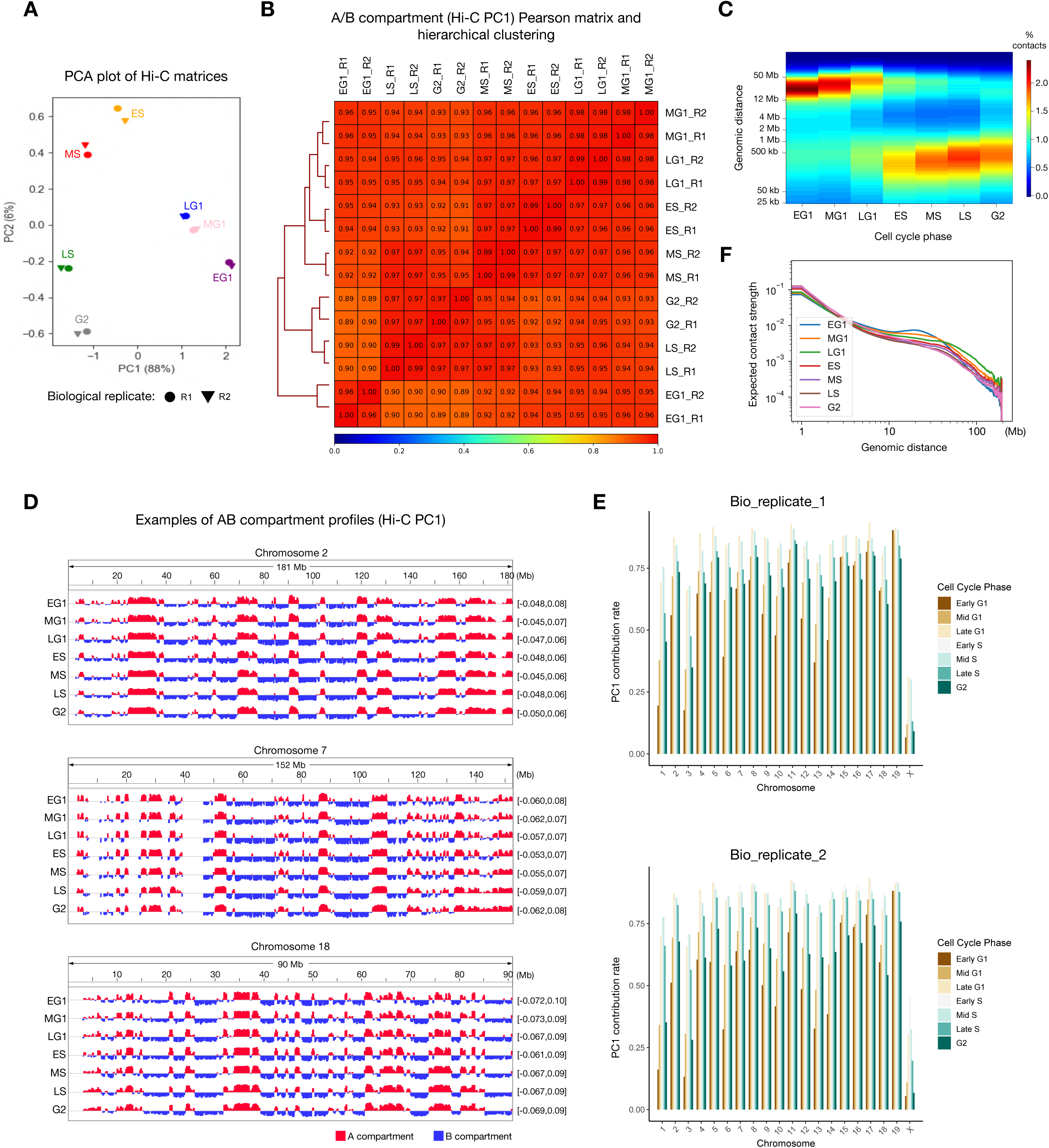
Quality control and analyses of Hi-C data from cell-cycle-phased mESCs. **(A)** Principal component analysis (PCA) of replicate Hi-C matrices. The plot shows PCA performed on Hi-C contact matrices (1-Mb resolution) from biological replicates of cell-cycle-phased samples from asynchronous populations, demonstrating replicate concordance. PCA sample size = 50,000. **(B)** Pearson correlation matrix and hierarchical clustering of Hi-C PC1 compartment profiles (1-Mb resolution) for biological replicates R1 and R2. **(C)** Contact decay profiles for all cell- cycle phases, plotted from 25 kb to 50 Mb, illustrating progressive changes in cis-interaction frequencies across genomic distances, including a gradual shift from long-range (> 12 Mb) to short-range (< 1 Mb) interactions during the -G1-to-S phase transition. **(D)** Representative IGV browser tracks of Hi-C PC1 compartment profiles (200-kb resolution) for chromosomes 2, 7, and 18. **(E)** Quantification of Hi-C PC1 contribution rates for individual chromosomes throughout interphase for replicate 1 (top) and replicate 2 (bottom) at 200-kb resolution. **(F)** Contact probability, P(s), plotted against genomic distance on a log-log scale (1-Mb resolution). Data shown in panels **(C)**, **(D)**, and **(F)** are from biological replicate 1 (representative of N=2 biological replicates). EG1, early G1; MG1, mid G1; LG1, late G1; ES, early S; MS, mid S; LS, late S.

**Figure S3.**
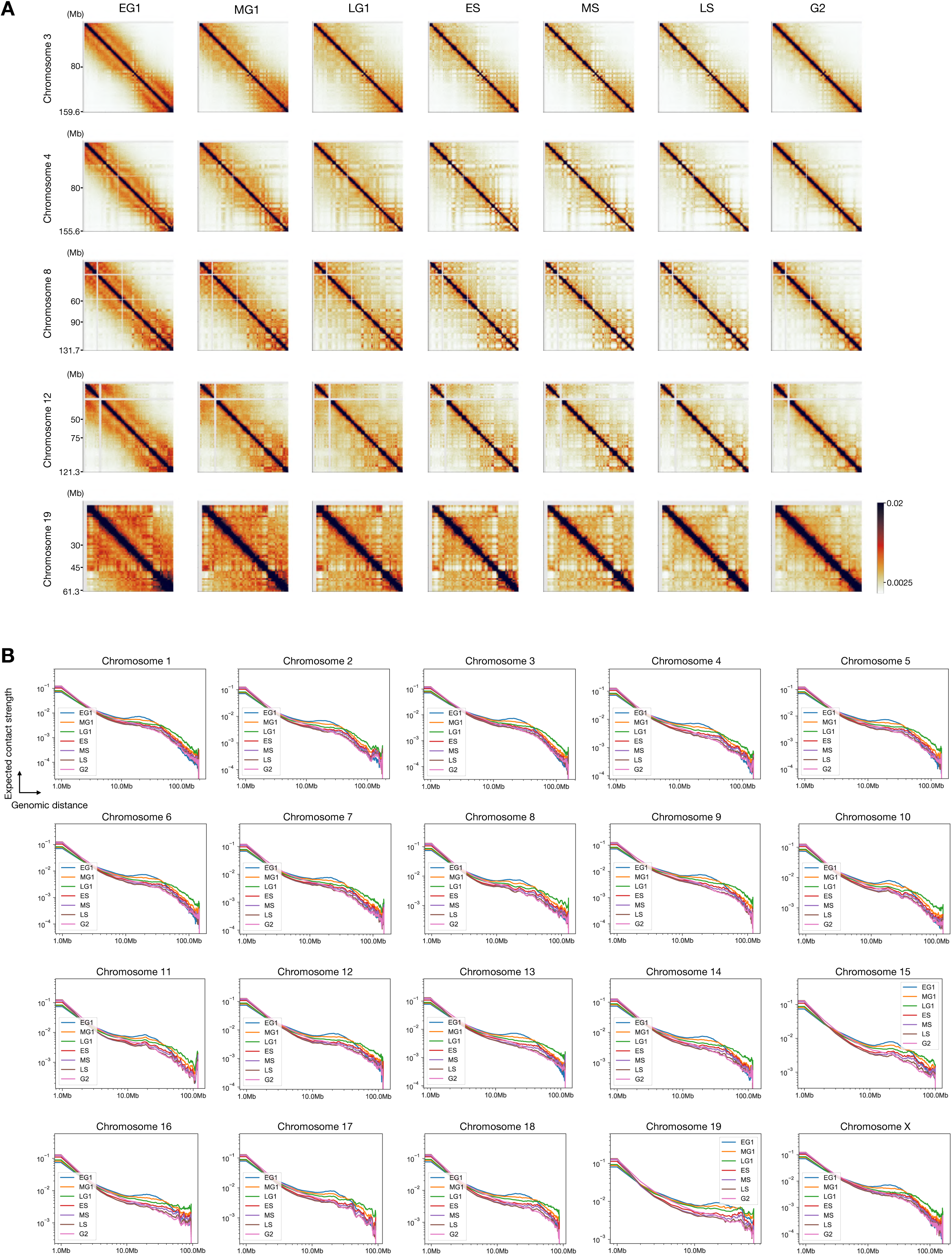
Cell-cycle-phased Hi-C data of individual chromosomes. **(A)** Hi-C contact maps (1-Mb resolution) of representative chromosomes (chr3, 4, 8, 12, and 19) across cell-cycle phases. **(B)** Contact probability, P(s), versus genomic distance on a log-log scale (1-Mb resolution) for individual chromosomes. Data are from merged biological replicates (N=2).

**Figure S4.**
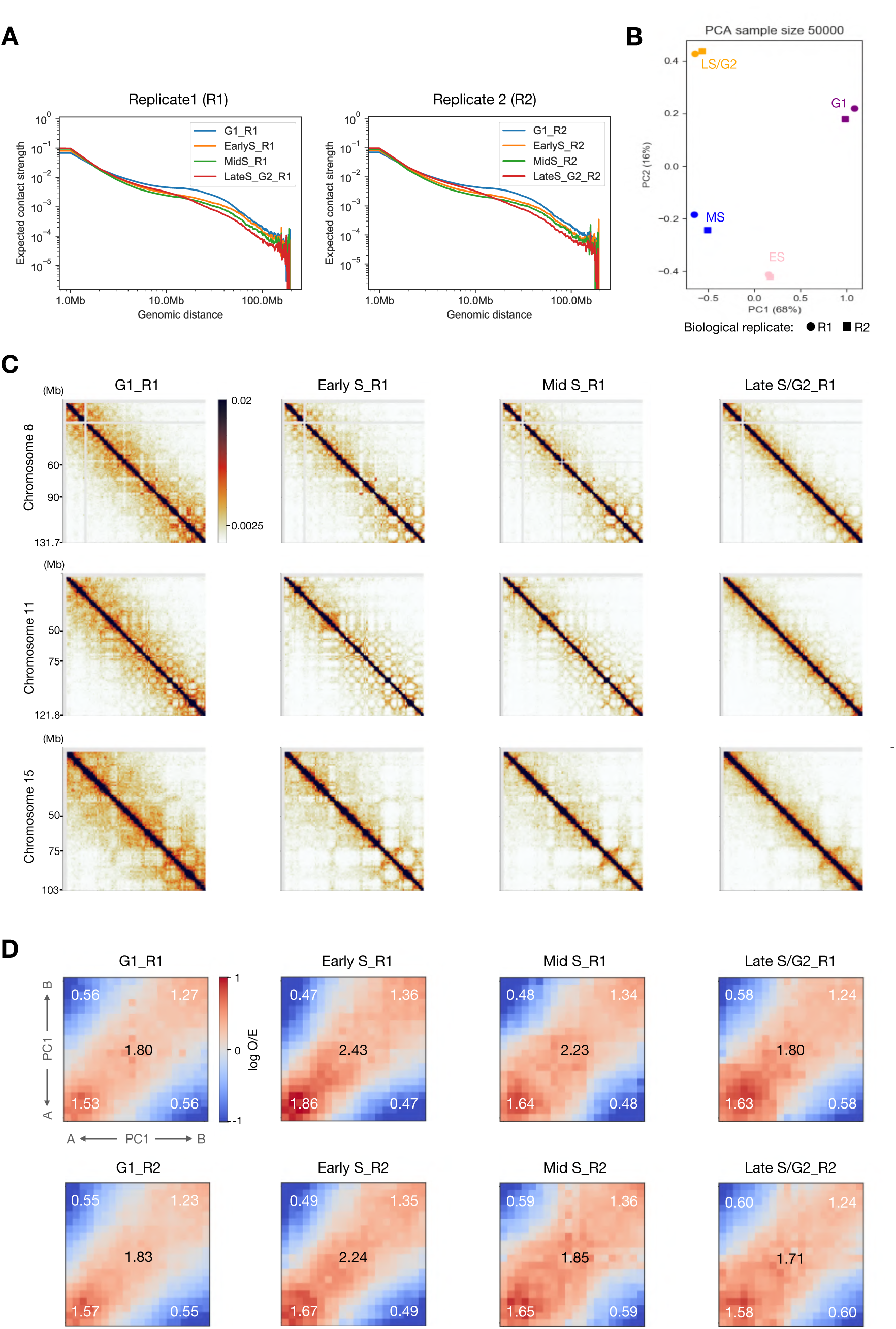
Re-analysis of single-cell Hi-C data from Nagano *et al.* ^12^ (pseudo-bulk analysis) **(A)** Contact probability versus genomic distance. Log-log plots of P(s) for merged single-cell Hi-C data (1-Mb resolution, N=2 biological replicates). Consistent with our findings, G1 phase exhibits the longest interaction range compared to mid- and late S/G2 phases. **(B)** Principal component analysis (PCA) of Hi-C contact matrices (1-Mb resolution; sample size = 50,000) demonstrates high reproducibility between biological replicates. **(C)** Hi-C contact maps (1-Mb resolution) of representative chromosomes (chr 8, 11, and 15) across cell-cycle phases of replicate 1. **(D)** Compartment strength quantification. Hi-C saddle plots for replicate 1 (top) and replicate 2 (bottom) at 1-Mb resolution show overall compartment strength (numeric values in black) and specific A-A, B-B, and A-B interaction frequencies (numeric values in white). The color scale represents observed/expected (O/E) contact frequencies in 5-percentile bins. As in our data, A/B compartment strength peaks during S-phase and diminishes in late S/G2.

**Figure S5.**
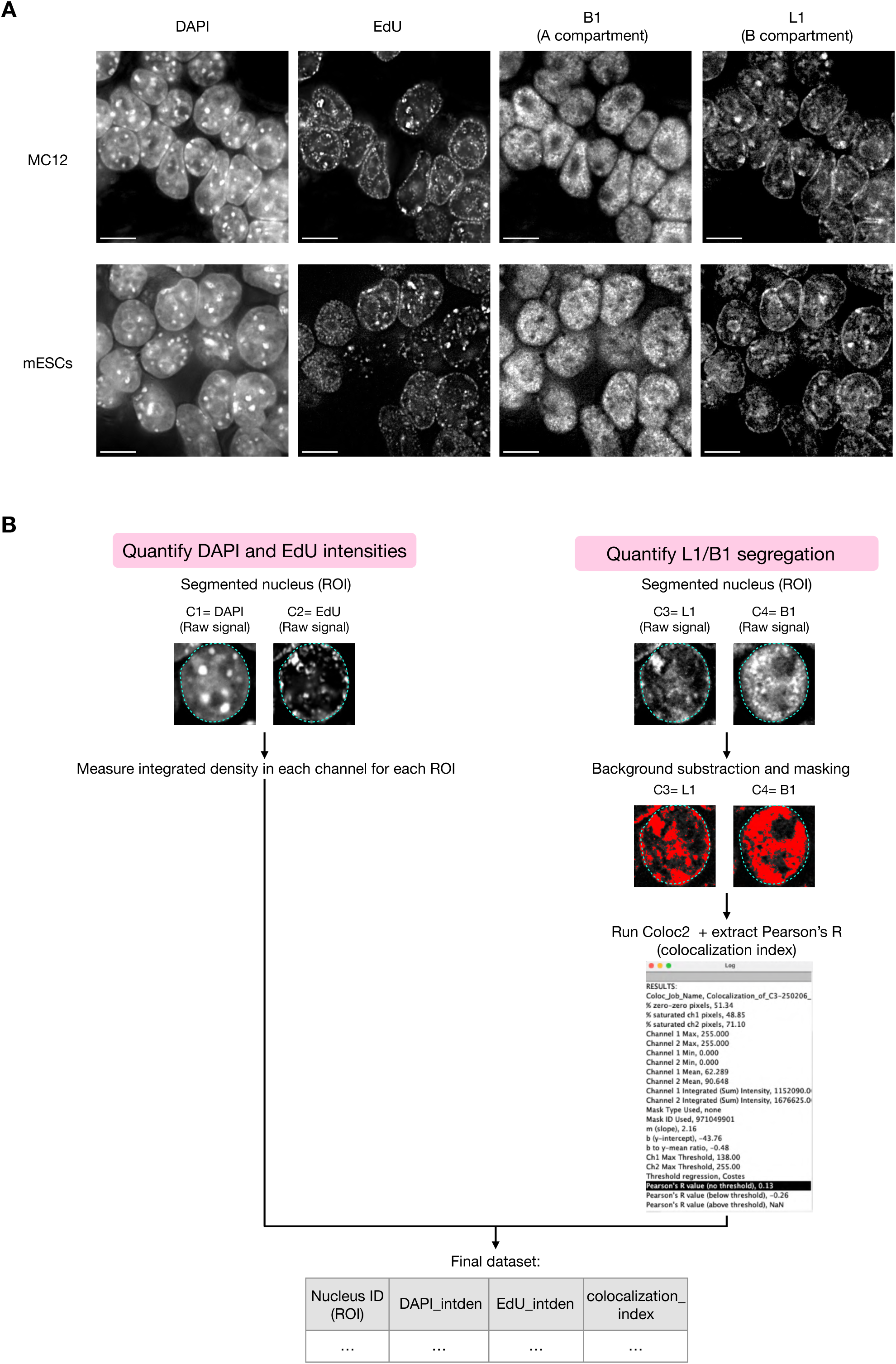
L1/B1 segregation analysis across the cell cycle in MC12 cells and mESCs. **(A)** Representative images of MC12 (top) and CBMS1 mESC (bottom) nuclei following L1/B1-EdU FISH. Scale bar= 10 µm. **(B)** Workflow for generating the dataset used for quantification of L1/B1 segregation across cell-cycle stages (see Methods for details). C, channel; ROI, region of interest.

**Figure S6.**
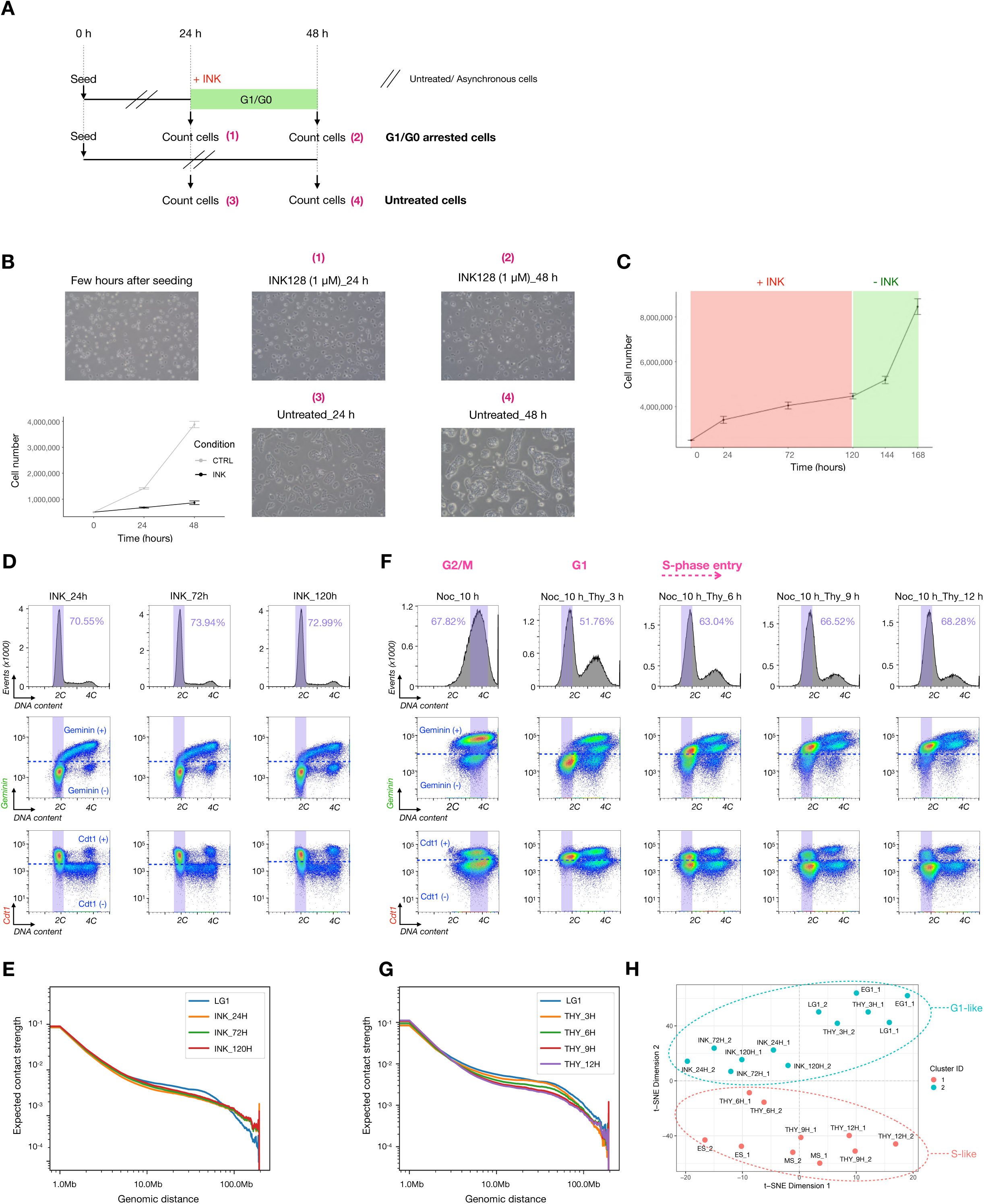
Validation of G1/G0 and G1/S arrest and comparison of Hi-C compartments in mESCs. **(A-C)** Reversible G1/G0 arrest of mESCs by INK-128. **(A)** Schematic of the time-course cell-cycle arrest experiment with INK-128 (1 µM). INK, INK-128. **(B)** Representative images of cells under each condition (numbered as in (A)), with corresponding cell-count quantification (mean ± SD, N=3). **(C)** Cell proliferation following long-term (120 h) INK-128 treatment and release into fresh medium (mean ± SD, N=3). SD, standard deviation. **(D)** FACS analysis of INK-128-treated (G1/G0-arrested) cells. DNA content (top panel), Geminin-mVenus (log scale) versus DNA content (middle panel), and Cdt1-mCherry (log scale) versus DNA content (bottom panel). Purple gates indicate cell populations sorted for Hi-C. **(E)** Contact probability, P(s), versus genomic distance for INK-128-arrested cells (log-log scale, 1-Mb resolution). Data are from single biological replicates of independent experiments (N=2). **(F)** FACS analysis of Nocodazole/Thymidine-treated (G1/S-arrested) cells. Panels are arranged as in (D). Cell-cycle stages (labeled in pink) were inferred for each population based on Geminin and Cdt1 fluorescence. **(G)** Contact probability, P(s), versus genomic distance for G1/S-arrested cells (log-log scale, 1-Mb resolution). Data are from single biological replicates of independent experiments (N=2). **(H)** t-SNE analysis with k-means clustering (k=2) of compartment saddle plot data (1-Mb resolution) from asynchronous cell-cycle samples (early G1, late G1, early S, mid S) and synchronized samples (INK-128-treated and Nocodazole + Thymidine-treated). Labels “1” and “2” denote biological replicates 1 and 2, respectively.

**Figure S7.**
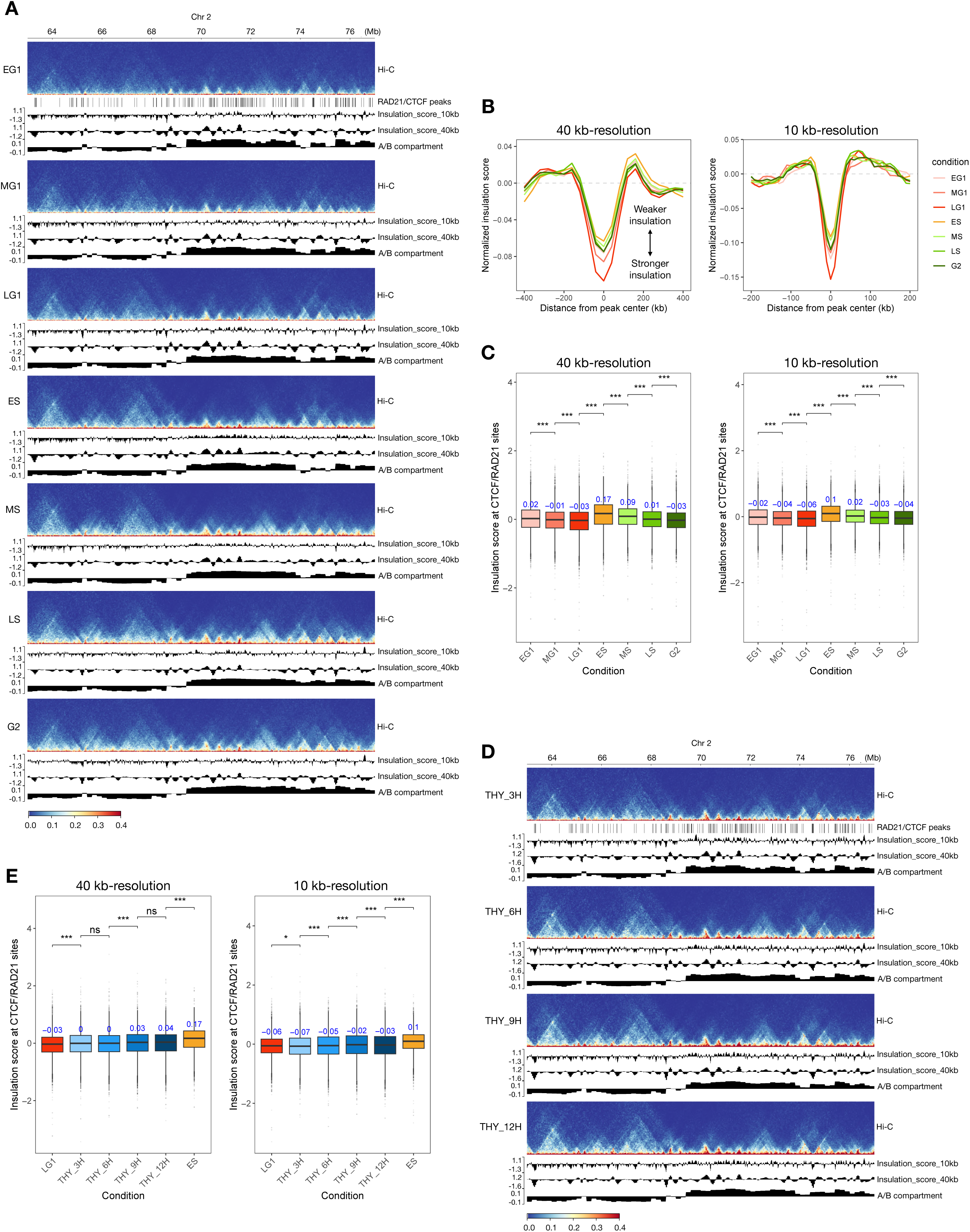
Insulation score dynamics at RAD21/CTCF sites during normal cell- cycle progression and upon G1/S arrest. **(A)** Genome browser snapshots of a representative region on chromosome 2 for the indicated cell cycle stages. RAD21/CTCF dual-occupancy sites are shown as a single track at the top (data from Hansen *et al*. 2017^26^). For each stage, the following tracks are displayed: Hi-C contact map (40- kb resolution), insulation scores at two resolutions (10- kb resolution using a 50- kb sliding windows and 40- kb resolution using 200-kb sliding windows), and A/B compartment track (200- kb resolution). **(B)** Meta-plots showing genome-wide average insulation scores centered at RAD21/CTCF sites (1000 randomly sampled peaks) for the indicated cell cycle stages. Insulation was calculated at two resolutions: left, 40- kb resolution using 200- kb sliding windows (±400 kb from peak center); right, 10- kb resolution using 50- kb sliding windows (±200 kb from peak center). **(C)** Boxplots showing genome-wide insulation scores at RAD21/CTCF dual-occupancy sites for the indicated cell- cycle stages. Median values are shown in blue. Pairwise comparisons between consecutive conditions were performed using the Wilcoxon rank-sum tests; ns, not significant; *, *P* ≤ 0.05; **, *P* ≤ 0.01; ***, *P* ≤ 0.001; ****, *P* ≤ 0.0001. **(D)** Same as (A) but showing data for G1/S-arrested cells at the indicated time points following release from nocodazole and thymidine treatment. **(E)** Boxplots as in (C), but comparing insulation scores between late G1, thymidine-arrested cells, and early S-phase cells. Data are from merged biological replicates (N=2).

**Figure S8.**
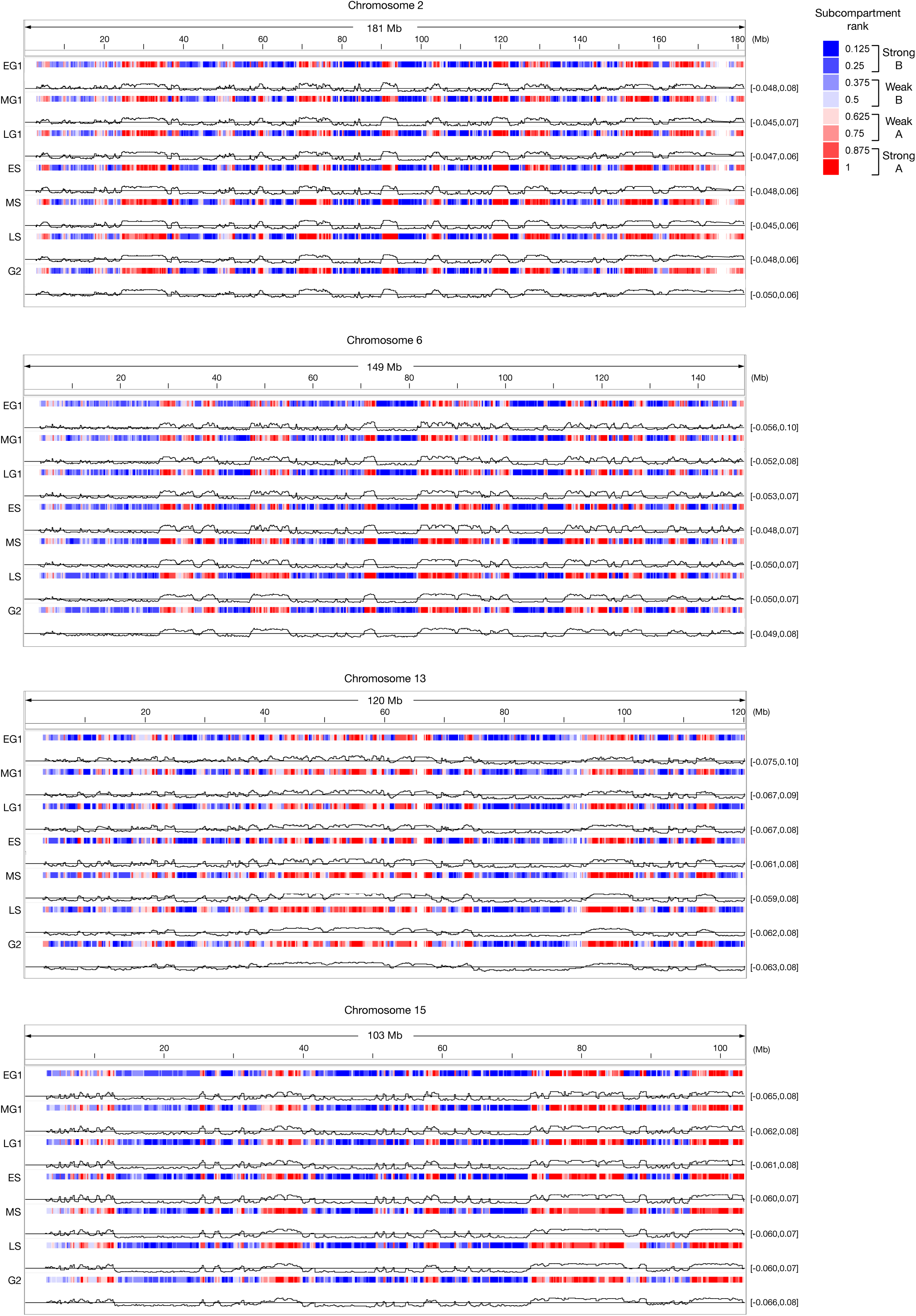
Calder subcompartment organization of representative chromosomes across interphase. IGV browser tracks of Calder subcompartments (40-kb resolution) for representative chromosomes (chr 2, 6, 13 and 15) across all cell-cycle stages. Data are from merged biological replicates (N=2).

**Figure S9.**
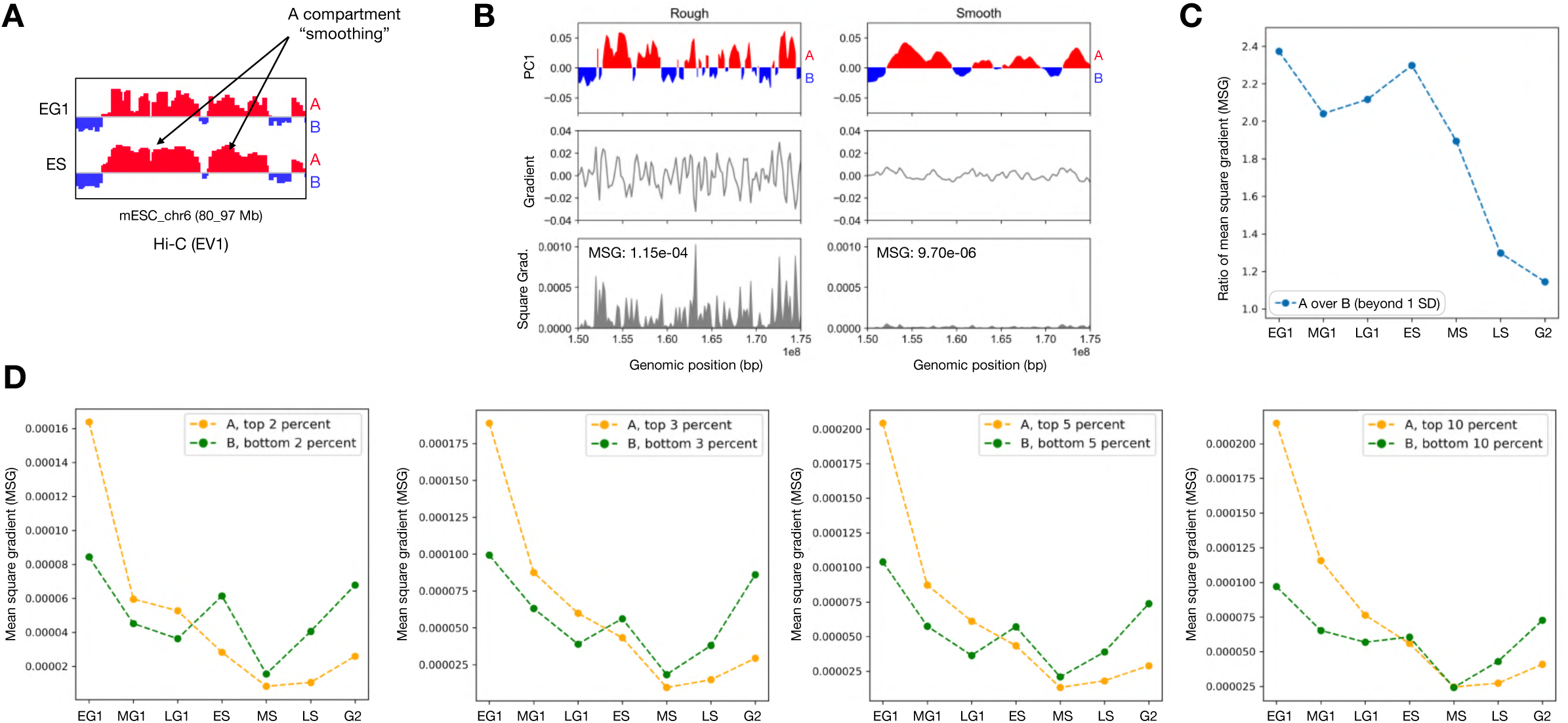
Quantification of A/B compartment PC1 signal “smoothness” by mean-square gradient (MSG) **(A)** Example genomic region showing Hi-C PC1 compartment profiles on IGV, illustrating smoothness of the A-compartment signal in S phase compared with G1. **(B)** (Top) Two schematic examples of compartment-like signals. The signal on the right is an artificially smoothed version of the signal on the left using a sliding-window average. (Middle) The gradient of each curve is plotted, revealing reduced variation in the smoothed curve, while keeping the same vertical limits. (Bottom) The squared gradient for each curve is plotted. The corresponding mean-square gradient (MSG) values are indicated. The smooth curve shows an MSG over tenfold lower, demonstrating the utility of MSG for quantifying smoothness. **(C**) Ratio of MSG between strong A (top > 1 SD; SD, standard deviation) and strong B (bottom > 1 SD) compartment signals (Hi-C PC1) for each cell-cycle stage. **(D)** MSG values for compartments defined by different thresholds. From left to right, compartments are defined with increasingly stringent thresholds: the top versus bottom 10%, 5%, 3%, and 2% of Hi-C PC1 values. Data points represent average MSG values. Data in **(C, D)** are from merged biological replicates (N=2), analyzed at 200-kb resolution.

**Figure S10.**
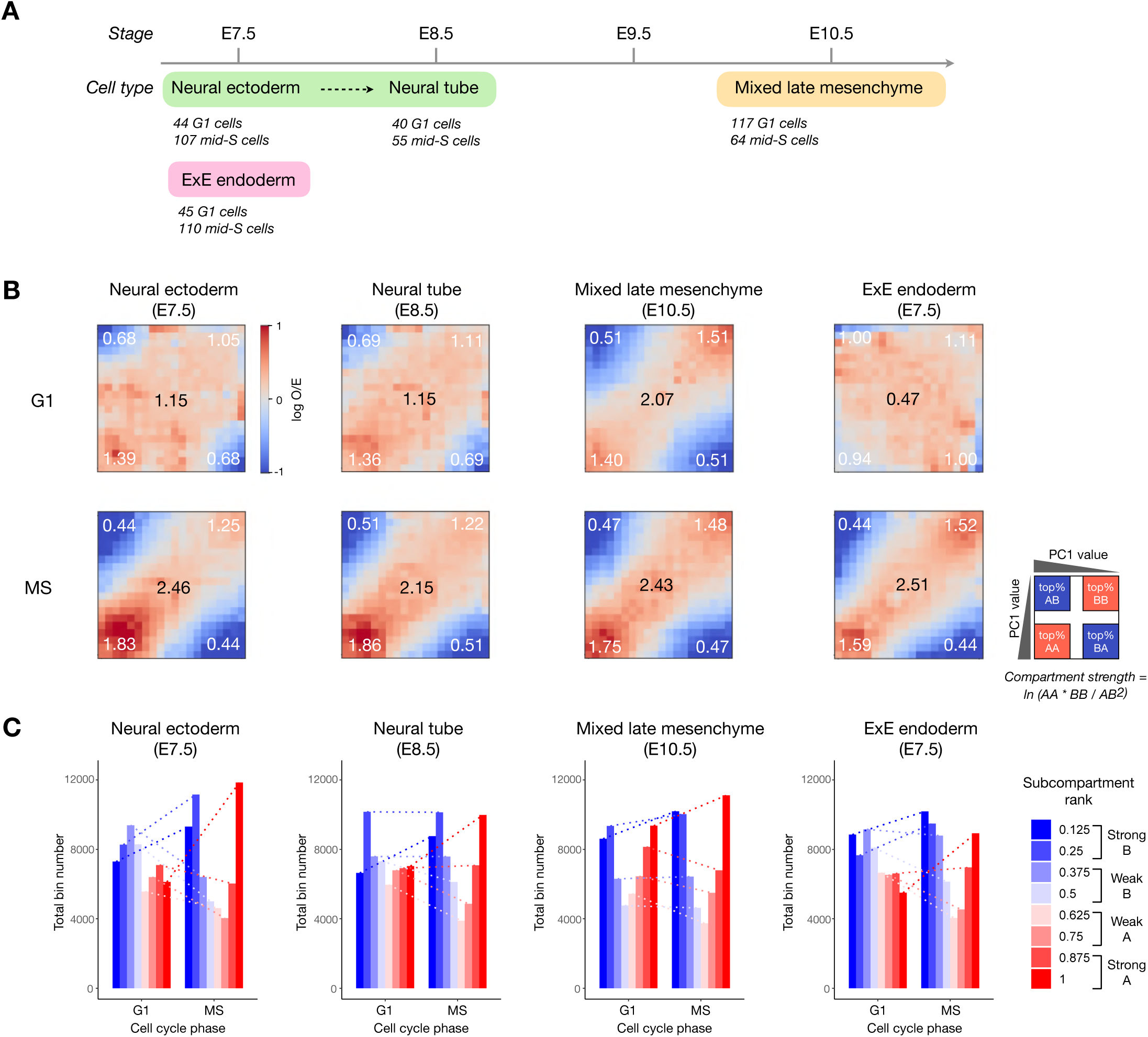
Pseudo-bulk Hi-C of single cells reveals conserved A-compartment consolidation during S phase across embryonic development. **(A)** Selection of single-cell populations from Liu *et al*.^29^ (HiRES) data for pseudo-bulk Hi-C analysis. The schematic shows the number of cells and developmental stage for each G1 and mid-S population. E7.5, embryonic day 7.5; ExE, extra-embryonic. **(B)** Hi-C saddle-plot analysis of compartment strength for data generated from the merged single-cell populations in (A) (1-Mb resolution), quantifying overall strength (numerical values in black) and specific AA, BB, and AB interaction frequencies (numerical values in white). The schematic on the right defines the axis ordering. The color scale represents O/E contact frequencies in 5-percentile increments. MS, mid S. **(C)** Abundance of Calder subcompartment ranks (200-kb resolution), shown as the total number of genomic bins, comparing G1 and mid-S phases across all four developmental stages.

**Figure S11.**
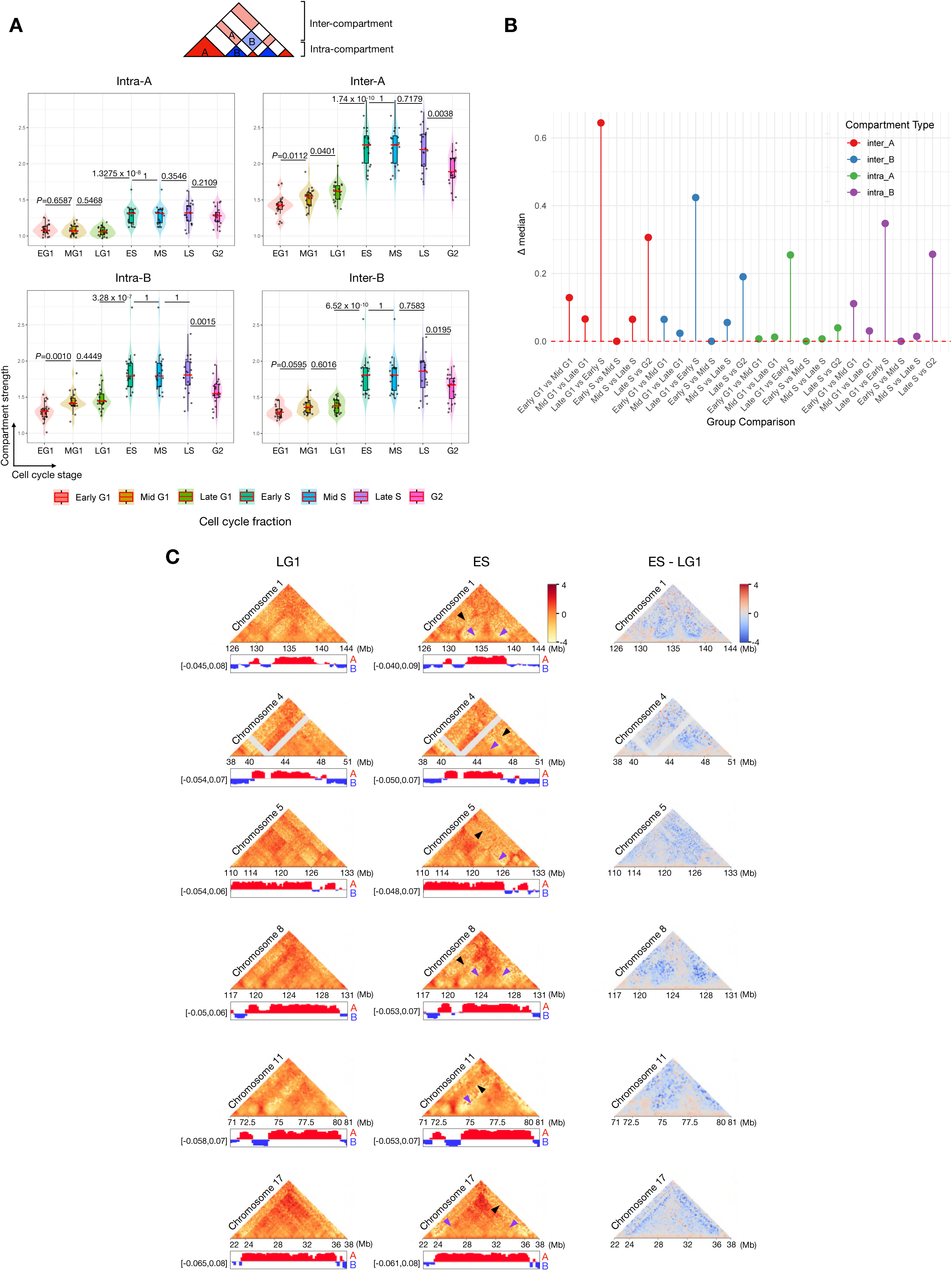
Representative genomic regions showing A-domain reorganization during S-phase. **(A)** Hi-C compartment strength for different interaction types (inter-A, intra-A, inter-B, intra-B) across interphase, from early G1 (EG1) to G2 phase. Distributions are shown as violin plots with medians indicated by red bars (each dot represents a single chromosome). Statistical significance between consecutive cell-cycle stages was assessed using pairwise Wilcoxon rank-sum tests. **(B)** Quantification of the Δ median interaction strength between consecutive cell-cycle stages for all interaction types. Inter-A compartment interactions show the largest increase during the late G1-to-early S-phase transition. **(C)** Observed/Expected Hi-C matrices and corresponding PC1 compartment profiles for six genomic regions (listed below), comparing late G1 (LG1, left panel) and early S phase (ES, middle panel). Purple arrowheads indicate loss of intra-A compartment signal in early S phase relative to late G1, while black arrowheads mark reduced contact frequency between neighboring A compartments. The right panel shows the differential contact heatmap (ES – LG1, early S – late G1). Region coordinates from top to bottom: chr1: 126–144 Mb; chr4: 38–51 Mb; chr5: 110–133 Mb; chr8: 117–131 Mb; chr11: 71–81 Mb; chr17: 22–38 Mb.

**Figure S12.**
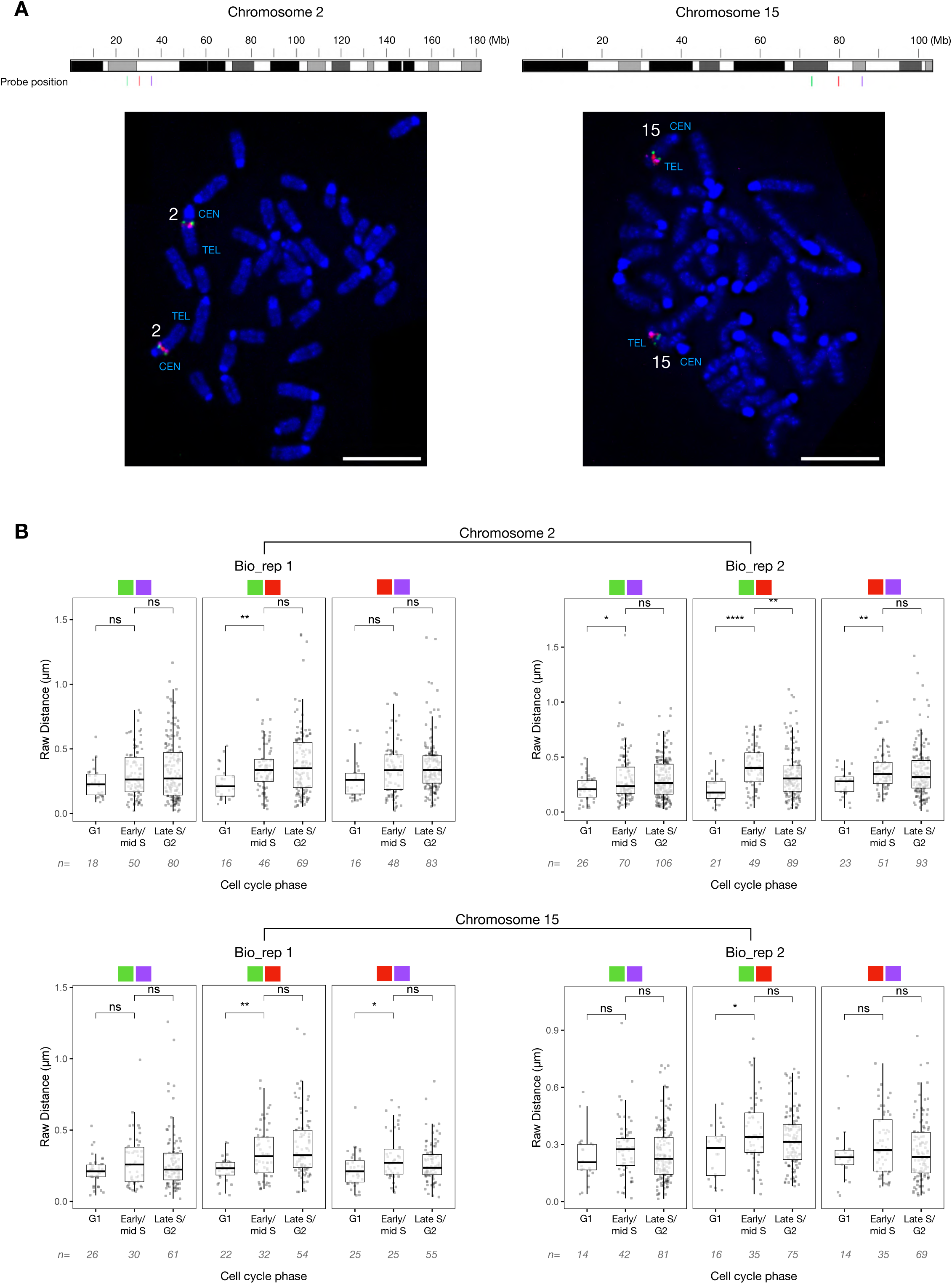
Validation of BAC probe positions and biological replicate analysis of inter-probe distances. **(A)** IGV browser view of BAC probe positions on the linear chromosome (top) and representative metaphase FISH on CBMS1 mESCs after Colcemid treatment (2 hours) (bottom). FISH signals confirm the expected relative localization of probes on chromosomes 2 (left) and 15 (right). Chromosome numbers are indicated on the metaphase spread. CEN, Centromere; TEL, Telomere. Scale bar = 10 µm. **(B)** Boxplots showing pairwise inter-probe distances (µm) separated by cell- cycle stage and presented for each biological replicate individually (N=2); n, number of nuclei. Top panels: chromosome 2; bottom panels: chromosome 15. For each chromosome, the three probe pairs are (from left to right): green–magenta, green–red, and red–magenta, as indicated above the boxplots. Pairwise comparisons between cell cycle stages (G1 vs early/mid S and early/mid S vs late S/G2) were performed using the Wilcoxon rank-sum test. P-values were adjusted for multiple comparisons using the Bonferroni method. ns, not significant; *, P ≤ 0.05; **, P ≤ 0.01; ***, P ≤ 0.001; ****P ≤ 0.0001.

**Figure S13.**
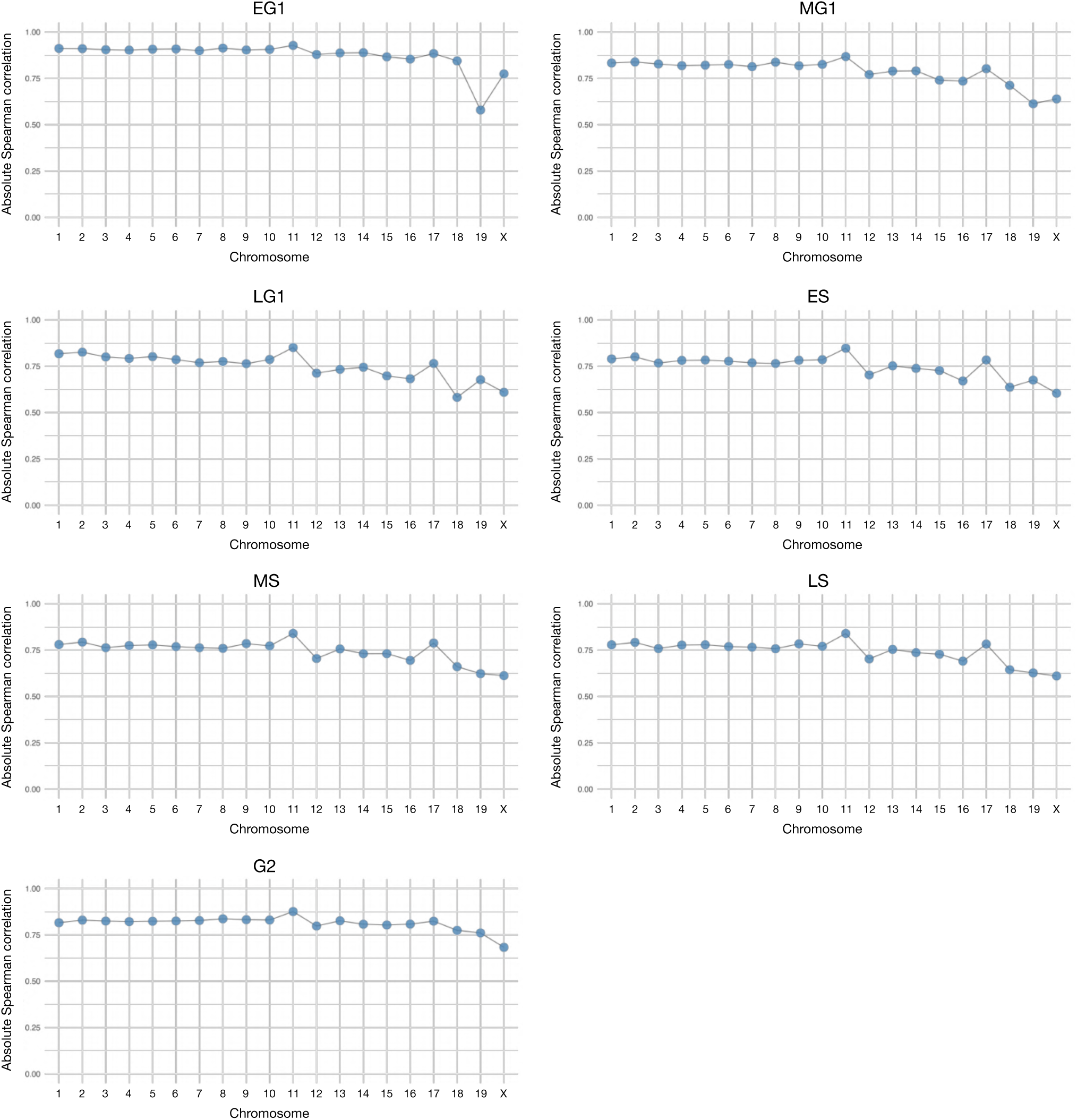
Correlation between reconstructed 3D structures and Hi-C data across the cell cycle. Absolute Spearman correlation coefficients between experimentally derived Hi-C contact matrices (200-kb bins) and distances from 3D genome structures reconstructed using the LorDG-3D Modeler in GenomeFlow^32^ using a conversion factor of 0.6 (10,000 iterations). Line plots show correlations for all chromosomes across different cell-cycle phases.

**Figure S14.**
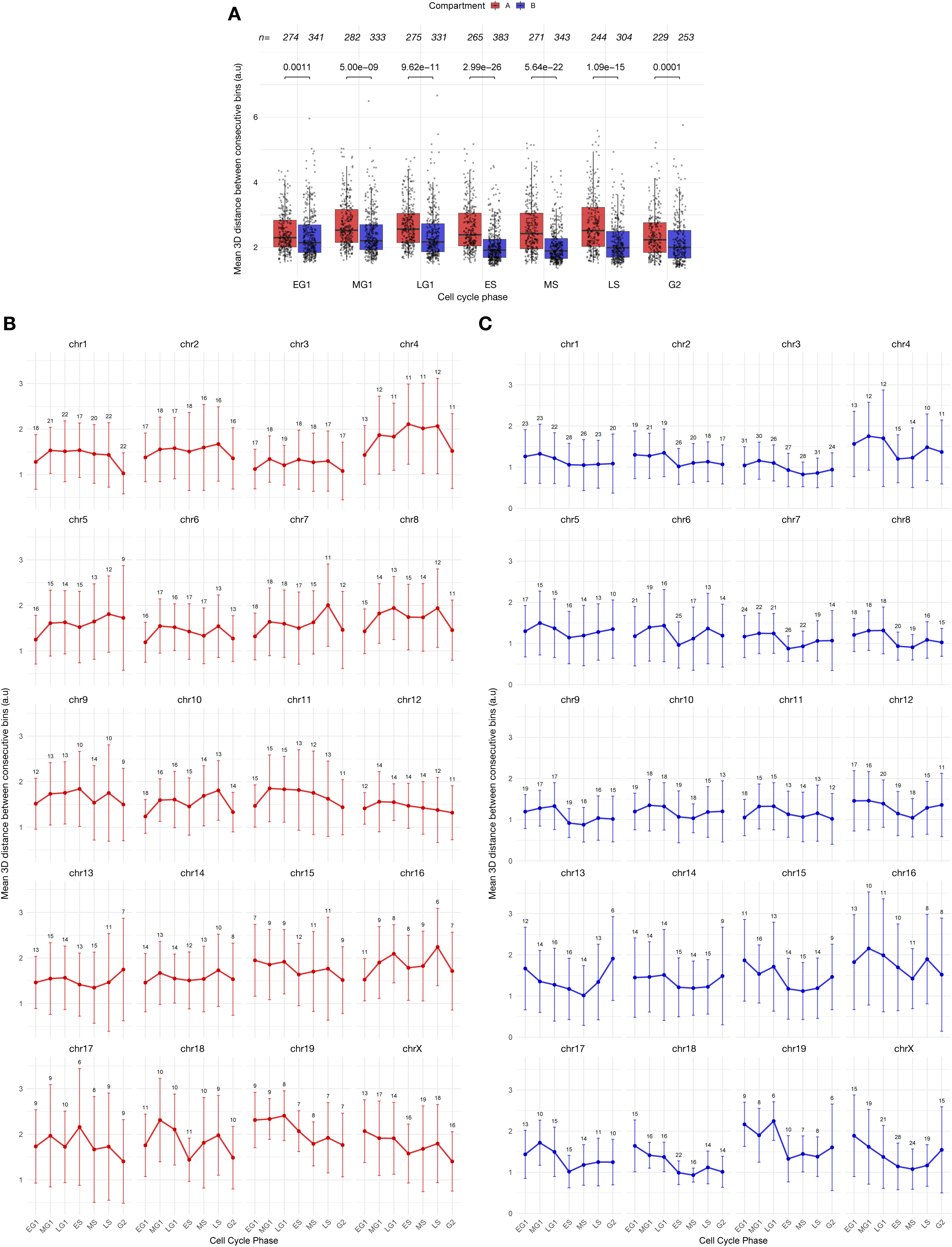
Simulated 3D dynamics of A and B compartment domains across cell-cycle phases. **(A)** Comparison of mean bin-to-bin distances between A and B compartment domains (> 1 Mb) across cell-cycle phases. Boxplots show the median and interquartile range. Each point represents a single domain. Pairwise comparisons between consecutive phases were performed using the Wilcoxon rank-sum test, and p-values were adjusted using the Benjamini-Hochberg method. **(B,C)** Mean bin-to-bin distances for A-compartment domains **(B)** and B-compartment domains **(C)** separated by chromosome. Data are presented as mean ± SD (points indicate means and error bars indicate SD). Numbers indicate the total number of domains per chromosome. Data in panels **(A–C)** are from merged biological replicates (N=2) analyzed at 200-kb resolution.

